# α-Synuclein and γ-Tubulin Cooperatively Regulate Activity-Evoked Presynaptic Microtubule Nucleation to Gate Dopamine Release

**DOI:** 10.64898/2026.05.29.728874

**Authors:** Alessandro Comincini, Sejoon Choi, Eugene V. Mosharov, Ryan Dosumu-Johnson, Edoardo Milanetti, Ellen Kanter, Giancarlo Ruocco, Graziella Cappelletti, David Sulzer, Francesca Bartolini

## Abstract

How α-synuclein regulates activity-dependent dopamine neurotransmission remains a longstanding question in synaptic biology. Here, we uncover a mechanism by which α-synuclein couples neuronal activity to interbouton synaptic vesicle (SV) transport and the local assembly of microtubules (MTs) at boutons. Using live imaging of dopaminergic neurons, we discover that neuronal activity triggers γ-tubulin-dependent MT nucleation preferentially at sites of SV fusion, promoting interbouton SV transport and striatal dopamine release. We find that α-synuclein cooperates with γ-tubulin to regulate this process through direct interactions with γ-tubulin and α/β-tubulin heterodimers. This function is gated by activity-dependent PLK2-mediated phosphorylation of α-synuclein at serine 129, which enhances tubulin binding and acts as a molecular switch for presynaptic MT nucleation. The Parkinson’s disease-associated α-synuclein mutations (A53T, A30P, and E46K) disrupt this regulation, uncoupling presynaptic MT dynamics from neuronal activity. Together, our findings reveal a physiological role for α-synuclein in orchestrating activity-dependent cytoskeletal remodeling at active synapses, providing a molecular link between presynaptic MT nucleation, SV redistribution, and dopamine neurotransmission that may be compromised in Parkinson’s disease.

**Graphical Abstract:** 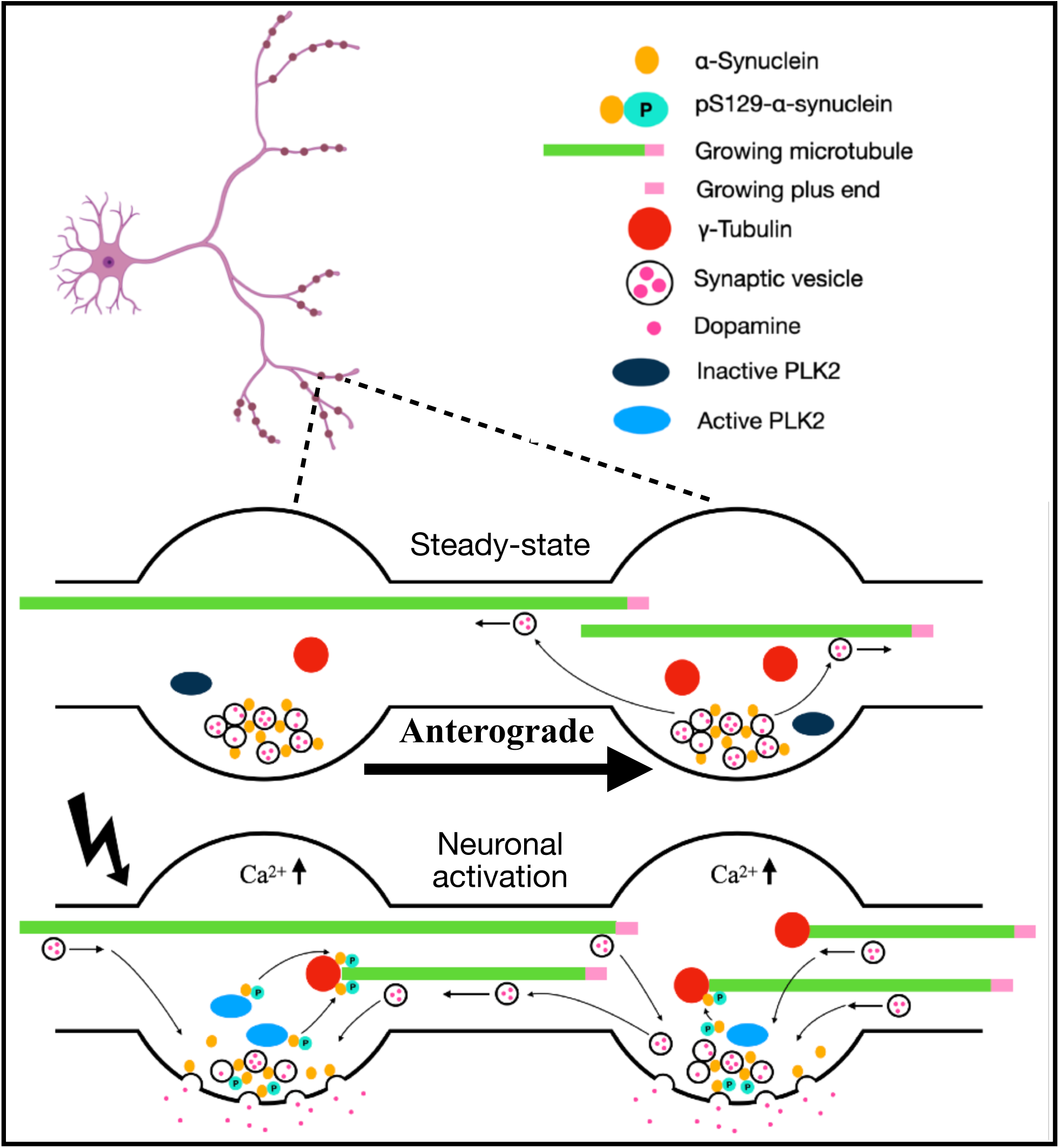

## INTRODUCTION

α-Synuclein is highly enriched at presynaptic terminals and has long been implicated in the regulation of neurotransmitter release, but the molecular mechanisms underlying its physiological function are incompletely understood (Oliveira et al., 2021; Sulzer & Edwards, 2019). In dopaminergic neurons, α-synuclein regulates activity-dependent dopamine release *in vitro* and *in vivo*. Early studies in coronal striatal slices revealed that α-synuclein loss impairs the replenishment of releasable dopamine during repeated stimulation (Abeliovich et al., 2000). More recent *in vivo* Fast-Scan Cyclic Voltammetry (FSCV) studies have demonstrated that α-synuclein contributes to both rapid facilitation and slower depression of evoked dopamine release, suggesting that its physiological function depends on neuronal activity and the temporal demands of neurotransmission (Shashaank et al., 2023; Somayaji et al., 2020). However, how α-synuclein couples neuronal activity to the mechanisms that sustain dopamine release remains unknown. This question is particularly relevant for dopaminergic neurons of the substantia nigra pars compacta (SNc), which possess exceptionally extensive axonal arbors, exhibit characteristic tonic and phasic firing patterns, and contain a substantial proportion of functionally silent presynaptic boutons that harbor SVs but do not release neurotransmitter (Gonon et al., 2000; C. Liu et al., 2018; Matsuda et al., 2009; Pereira et al., 2016; Rice & Cragg, 2008). These distinctive properties place considerable demands on the mechanisms that distribute and replenish SVs across axonal dopamine release sites. Dopaminergic neurons are also particularly vulnerable to α-synuclein pathology in Parkinson’s disease (PD), where synaptic dysfunction can precede overt neurodegeneration. Although the toxic effects of aggregated α-synuclein have received considerable attention, disruption of its normal presynaptic functions may also contribute to dopaminergic vulnerability. Understanding how α-synuclein supports neurotransmission is therefore essential for defining its physiological role and its potential contribution to disease.

MTs are emerging as important regulators of presynaptic function. Beyond their established roles in maintaining neuronal architecture and supporting long-range axonal transport, dynamic MTs undergo continuous cycles of growth and shrinkage and contribute to synaptic organization, plasticity, and neurotransmitter release (Baas et al., 2016; Dent, 2020; Parato & Bartolini, 2021; Waites et al., 2021). Neuronal MT arrays are maintained through spatially restricted, non-centrosomal nucleation mechanisms that enable local cytoskeletal remodeling within distinct subcellular compartments (Rolls, 2022; Sánchez-Huertas et al., 2016; Yogev et al., 2016). At presynaptic sites, dynamic MTs regulate SV cycles and provide tracks for the delivery of SV precursors and active zone components (Aiken & Holzbaur, 2024; Babu et al., 2020; Guedes-Dias et al., 2019; Guillaud et al., 2017). Importantly, neuronal activity stimulates γ-tubulin-dependent MT nucleation at *en passant* boutons in hippocampal neurons, providing a mechanism for the rapid generation of local transport tracks during synaptic activity (Qu et al., 2019). Whether a similar mechanism operates in dopaminergic neurons, whether MT nucleation is preferentially associated with functionally active release sites, and how neuronal activity engages the molecular machinery responsible for this process remain unknown.

A potential link between presynaptic MT dynamics and dopamine neurotransmission is provided by the ability of α-synuclein to interact directly with tubulin. Biochemical studies have shown that α-synuclein binds to soluble tubulin, undergoes tubulin-induced conformational changes, and promotes MT nucleation and dynamics *in vitro*, leading to its proposed function as an MT "dynamase" (Cartelli et al., 2016). α-Synuclein also associates with tubulin at striatal dopaminergic presynaptic terminals (Amadeo et al., 2021; Cartelli et al., 2016; Zhou et al., 2010). Notably, the PD-associated α-synuclein variants A30P, E46K, and A53T alter tubulin-induced folding and MT assembly *in vitro*, suggesting that disruption of α-synuclein–tubulin interactions may affect neuronal MT organization (Cartelli et al., 2016). However, whether these biochemical interactions regulate MT nucleation at presynaptic boutons, and whether this function contributes to dopamine neurotransmission, has not been established.

An intriguing possibility is that neuronal activity regulates the interaction of α-synuclein with the MT nucleation machinery through post-translational modification. Neuronal activation promotes phosphorylation of α-synuclein at serine 129 (S129), primarily through polo-like kinase 2 (PLK2), a modification that regulates its synaptic localization and interactions with presynaptic proteins (Parra-Rivas et al., 2023; Ramalingam et al., 2023; Stavsky et al., 2024). Although S129 phosphorylation is strongly associated with pathological α-synuclein accumulation in synucleinopathies, emerging evidence supports an important physiological role for this modification in activity-dependent synaptic function. Whether S129 phosphorylation controls α-synuclein interactions with tubulin and thereby couples neuronal activity to local MT nucleation is unexplored.

Here, we uncover an activity-dependent presynaptic mechanism through which α-synuclein and γ-tubulin regulate MT nucleation in dopaminergic neurons. By combining live imaging of MT dynamics and SV trafficking with measurements of synaptic exocytosis and striatal dopamine release, we demonstrate that neuronal activity promotes γ-tubulin-dependent MT nucleation at *en passant* axonal boutons that are associated with active sites of SV release. We identify α-synuclein as a key regulator of this process and establish PLK2-dependent S129 phosphorylation as a molecular switch linking neuronal activity to tubulin binding and presynaptic MT initiation. Furthermore, we show that PD-associated α-synuclein mutations interfere with the normal coupling between neuronal activity and presynaptic MT nucleation. Together, our findings reveal a previously unrecognized physiological function of α-synuclein in linking synaptic activity to local cytoskeletal remodeling, and provide a molecular mechanism connecting presynaptic MT nucleation to SV redistribution and the kinetics of activity-dependent dopamine neurotransmission, and how that may be compromised in synucleinopathies.

## RESULTS

### Activity-induced microtubule initiation at en passant boutons is a conserved feature of dopaminergic neurons

Dopaminergic neurons of the SNc that project to the striatum are distinguished by extensive axonal arborization and tonic activity interrupted by episodes of phasic burst firing (Matsuda et al., 2009). While neuronal firing has been shown to increase MT dynamics in hippocampal neurons (Qu et al., 2019), the organization and regulation of dynamic MTs in dopaminergic neurons are unexplored. To examine these features, we expressed a FLEX version of the MT plus-end marker EB3-mGreenLantern in primary DAT-Cre mouse dopaminergic neurons from the ventral midbrain and monitored growing MT plus ends (comets) by live imaging. Following imaging, neurons were fixed and stained for the axon initial segment marker ankyrin G to distinguish axons from dendrites (**Fig. 1A, B**). The imaged neurites were subsequently identified, and their proximal (<100 µm) and distal (>100 µm) segments analyzed.

**Figure 1.**
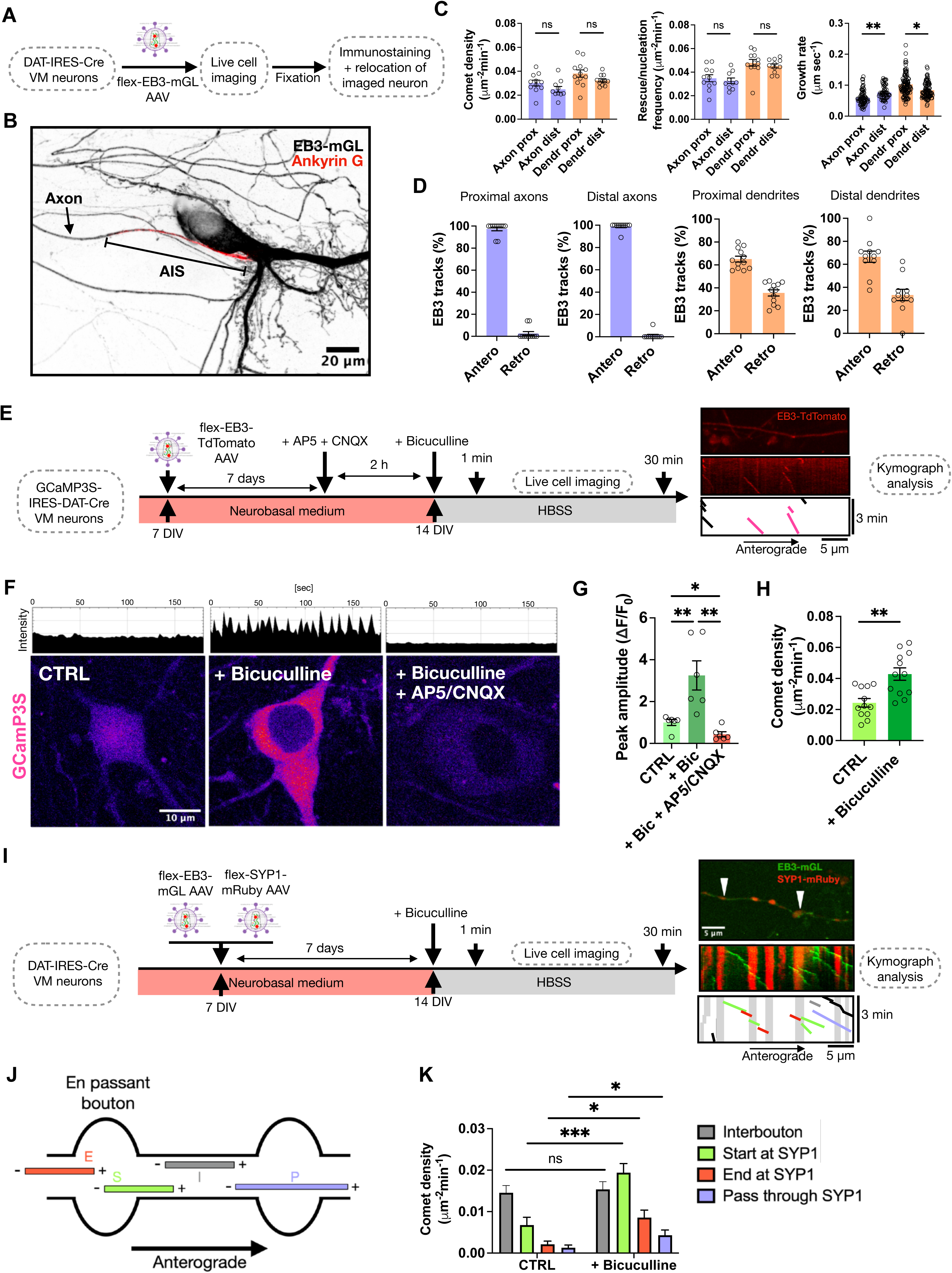
Activity-evoked dynamic microtubule initiation at *en passant* bouton is conserved in dopaminergic neurons and depends on glutamatergic stimulation. **(A)** Schematic of the experimental design. **(B)** Representative image of a DAT-Cre dopaminergic neuron fixed after live imaging and stained for Ankyrin G to enable post hoc identification of the axon initial segment. **(C)** Quantification of comet density, rescue/nucleation frequency, and growth rate of EB3 comets in proximal (x<100 µm from cell body) and distal (x>100 µm from cell body) axons and dendrites in DAT-Cre neurons (14-16 DIV). Data are mean ± SEM; ns, p > 0.05; Kruskal-Wallis non-parametric ANOVA. **(D)** Percentage of anterograde and retrograde EB3 comets among total EB3 tracks in proximal (x<100 µm from cell body) and distal (x>100 µm from cell body) axons and dendrites in DAT-Cre neurons (14-16 DIV). **(E)** Schematic of the experimental design with a representative frame and kymograph from live imaging of GCamP3S-DAT-IRES-Cre neurons expressing EB3-TdTomato. In the kymograph schematic, EB3 tracks are shown in black (incomplete) and magenta (complete). **(F)** Representative time-lapse images and whole-field fluorescence intensity profile from GCamP3S-DAT-IRES-Cre neurons upon bicuculline stimulation or NMDA/ AMPA receptors blockers to inhibit glutamatergic stimulation. **(G)** Peak GCamP3S fluorescence amplitude in GCamP3S-DAT-IRES-Cre neurons treated as in (F). Data are mean ± SEM; ** p<0.01, Kruskal-Wallis non-parametric ANOVA. **(H)** Quantification of comet density in GCamP3S-DAT-IRES-Cre upon bicuculline-induced neuronal activation. **(I)** Schematic of the experimental design with representative live-imaging frames and kymograph from distal axons of DAT-Cre neurons expressing flexed EB3-mGreenLantern and synaptophysin1 (SYP1)-mRuby. In the kymograph schematic, EB3 tracks are shown in black (incomplete), grey (interbouton), green (start at bouton), red (end at bouton), and purple (pass through bouton). **(J)** Schematic representation of the EB3 subclassification (interbouton, start, end, pass through) and color code as in (I) relative to stable SYP1^+^ puncta in axons of DAT-Cre neurons. **(K)** Quantification of subclassified EB3 (interbouton, start, end, pass through) comet density relative to stable SYP1^+^ puncta in axons of DAT-Cre neurons treated as in (I).

As seen in hippocampal neurons (Kollins et al., 2009; Qu et al., 2019), dopaminergic axons showed lower MT growth rates, comet density, and rescue/nucleation frequency than dendrites, which contained a larger fraction of highly dynamic MTs, with only minor differences between proximal and distal regions and no change in catastrophe frequency (**Fig. 1C, S1A**). MT polarity followed the canonical organization described in vertebrate neurons, with axons being almost entirely plus-end-out MTs and dendrites exhibiting mixed polarity (Baas et al., 1988; Yau et al., 2016) with approximately 65% of comets moving anterogradely and 35% retrogradely in both proximal and distal compartments (**Fig. 1D)**. These data indicate that, despite their distinctive firing properties and extensive axonal arborization, dopaminergic neurons maintain a canonical organization of MT polarity, with nearly all axonal EB3 comets moving anterogradely, consistent with predominantly plus-end-out MT polarity that support directional transport.

To assess whether presynaptic MT initiation in dopaminergic neurons is activity-dependent, neuronal activity was chemically induced by application of the GABA_A_ receptor antagonist bicuculline (**Fig. 1E, S1B**). This rapidly increased neuronal activity, as demonstrated by waves of increased cytosolic calcium that were completely abolished by co-application of NMDA and AMPA receptor antagonists, indicating that this induced calcium response depends on glutamatergic transmission (**Fig. 1F, G**). Importantly, bicuculline-induced activity increased total MT comet density by nearly two-fold (**Fig. 1H**), closely mirroring effects previously reported in excitatory hippocampal neurons (Qu et al., 2019). These findings demonstrate that increased neuronal activity enhances MT dynamics in dopaminergic axons.

To determine whether this activity-dependent increase in growing MT plus ends preferentially originated at presynaptic sites, synaptophysin1 (SYP1) was co-expressed to label SV clusters and presynaptic boutons (**Fig. 1I**). EB3 comets were classified according to their spatial relationship with stable presynaptic puncta. Remarkably, the activity-dependent increase in total comet density was entirely attributable to bouton-associated MT plus ends, including comets initiating at, terminating within, or passing through boutons (**Fig. 1J, K**), whereas interbouton comet density remained unchanged. MT initiation events at *en passant* boutons (starting comets) accounted for the largest proportion of this increase (**Fig. 1K**). Together, these data identify presynaptic boutons as the principal sites of activity-evoked MT initiation in dopaminergic axons, revealing a conserved mechanism by which neuronal activity drives spatially restricted cytoskeletal remodeling at presynaptic terminals.

### γ-Tubulin controls presynaptic microtubule nucleation in dopaminergic neurons

Because EB3 comet analysis cannot distinguish the rescue of pre-existing MTs from *de novo* nucleation, we first asked whether γ-tubulin, the core component of the MT nucleation machinery in developing axons and at *en passant* boutons of pyramidal neurons (Qu et al., 2019; Sánchez-Huertas et al., 2016), was also present in dopaminergic axons and positioned to mediate activity-induced MT initiation at presynaptic boutons. Immunostaining of dopaminergic neuronal cultures at 16 days *in vitro* (DIV) revealed enrichment of γ-tubulin signal within presynaptic regions of dopaminergic axons (**Fig. 2A**). Pixel-based colocalization analysis within a tyrosine hydroxylase (TH) mask to restrict the analysis to dopaminergic neurons showed that although only ∼10% of synapsin1/2 signal overlapped with γ-tubulin, ∼60% of γ-tubulin signal overlapped with synapsin1/2 (**Fig. 2B**). These values reflect intensity-based overlap and suggest that γ-tubulin is preferentially associated with presynaptic regions, potentially identifying boutons competent for MT nucleation.

**Figure 2.**
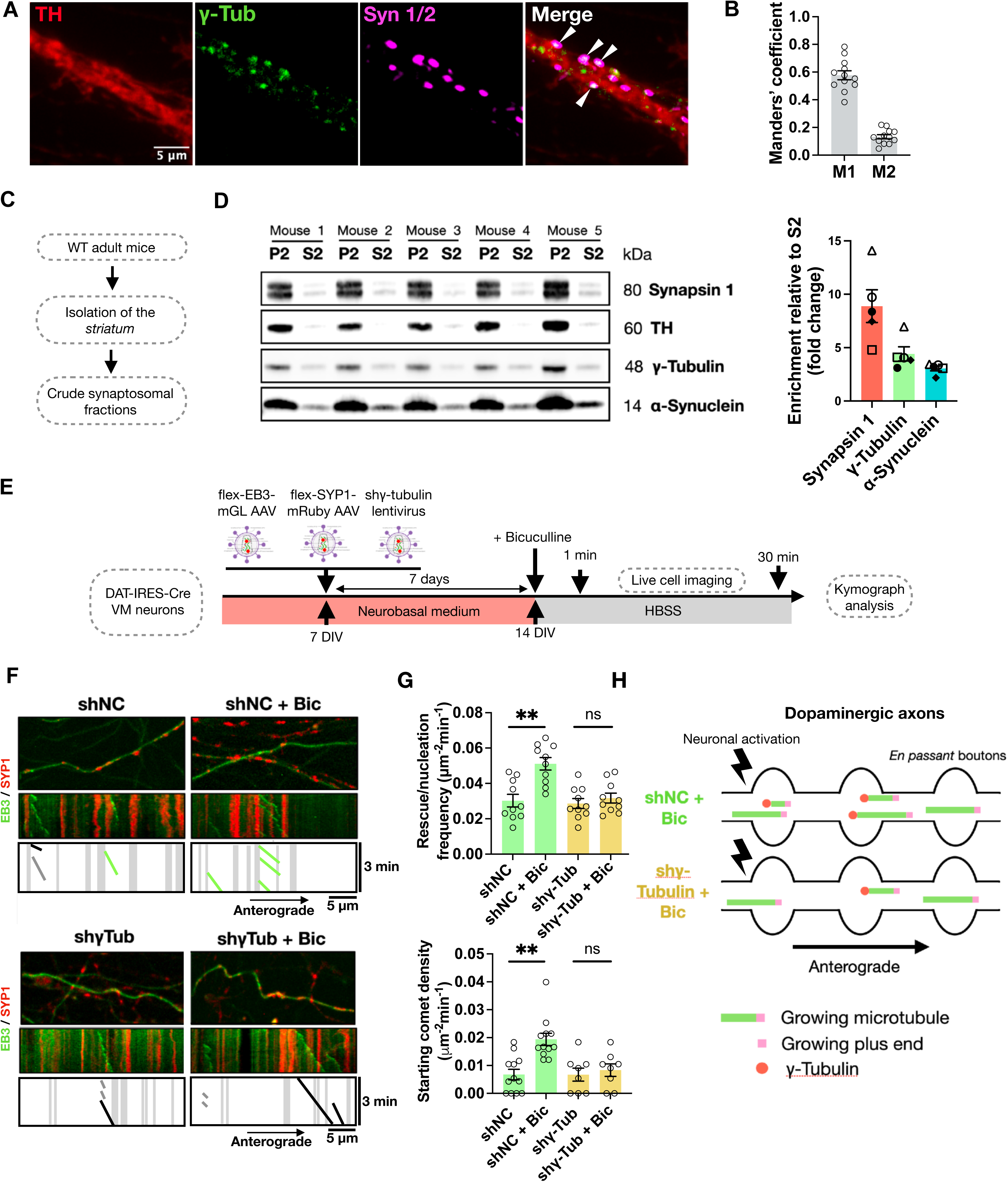
γ-Tubulin controls presynaptic microtubule nucleation in dopaminergic neurons. **(A)** Maximum-intensity projection of spinning disk confocal images from wild-type ventral midbrain neurons (16 DIV) fixed and stained for γ-tubulin, TH, and synapsin 1/2. White arrows indicate γ-tubulin puncta co-localizing with synapsin 1/2 within a TH mask. **(B)** Manders’ colocalization coefficients for synapsin1/2 and γ-tubulin in dopaminergic neurons treated as in (A). M1 denotes the fraction of γ-tubulin overlapping with synapsin1/2 within the TH mask; M2 denotes the fraction of synapsin1/2^+^ puncta overlapping with γ-tubulin within the TH mask. **(C)** Schematic of the protocol used to isolate and purify crude synaptosomal fractions from the mouse corpus striatum. **(D**) Crude synaptosomal fractions (P2) and corresponding supernatant fractions (S2) from the striatum of five 3-month-old WT mice, with quantitative western blot analysis of synapsin1, γ-tubulin, and α-synuclein. Bands were normalized to total protein (Fig. S2) and expressed as a fold change relative to S2. **(E)** Schematic of the experimental design for live-imaging in DAT-Cre neurons. **(F)** Representative kymographs from distal axons of DAT-Cre neurons expressing flexed EB3-mGreenLantern and SYP1-mRuby after 7 days of lentiviral delivery of shRNA targeting γ-tubulin (shγ-Tub) or non-coding control (shNC), at baseline and after bicuculline treatment (shNC + Bic and shγ-Tub + Bic). **(G)** Quantification of rescue/nucleation frequency of EB3 comets in distal (x>100 µm from cell body) axons and starting comet density relative to stable SYP1^+^ puncta in axons of DAT-Cre neurons treated as in (F). Data are mean ± SEM; ** p<0.01, Kruskal-Wallis non-parametric ANOVA. **(H)** Schematic representation of the summary of γ-tubulin knockdown on activity-evoked presynaptic MT nucleation in DAT-Cre neurons.

To independently validate this presynaptic localization, we isolated crude synaptosomal fractions from the corpus striatum (**Fig. 2C**), a region densely innervated by ventral midbrain dopaminergic axons. We found that γ-tubulin was enriched in the synaptosomal fraction, with levels approximately four-fold higher in P2 than in S2 (**Fig. 2D**), with minimal Golgi contamination (**Fig. S2B**). To test whether MT initiation at dopaminergic synapses depends on γ-tubulin, we depleted γ-tubulin using lentiviral shRNA (Qu et al., 2019) and quantified MT dynamics relative to stable presynaptic puncta (**Fig. 2E, S2C**). At baseline, γ-tubulin depletion did not affect total comet density or rescue/nucleation frequency in axons (shNC vs shγ-Tub). In contrast, upon bicuculline-induced neuronal activity, γ-tubulin knockdown completely abolished the activity-dependent increase in comet density observed in controls (shNC + Bic vs shγ-Tub + Bic) and selectively eliminated the activity-dependent increase in comets initiating at boutons (**Fig. 2F, 2G**) while sparing other bouton-associated events (**Fig. S2D**). Together, these results indicate that γ-tubulin is responsible for the activity-evoked increase in presynaptic MT initiation, supporting *de novo* nucleation as a major source of newly growing MTs in dopaminergic axons (**Fig. 2H**).

### Presynaptic microtubule nucleation promotes interbouton synaptic vesicle motility, and supports evoked dopamine release

Because presynaptic MT nucleation was selectively enhanced by neuronal activation, and dynamic MTs provide tracks for cargo transport along axons and between *en passant* boutons, we hypothesized that this mechanism might regulate dopaminergic neurotransmission. Dopaminergic neurons exhibit cell-autonomous tonic activity, including in culture as demonstrated above (**Fig. 1E-H**), so we first examined whether inhibiting γ-tubulin altered calcium activity. Incubation with the γ-tubulin inhibitor GatastatinG2 during bicuculline treatment did not change the amplitude of calcium transients measured by GCaMP3S fluorescence (**Fig. 3A, B**), indicating that γ-tubulin inhibition does not detectably alter activity-evoked calcium signaling in dopaminergic neurons.

**Figure 3.**
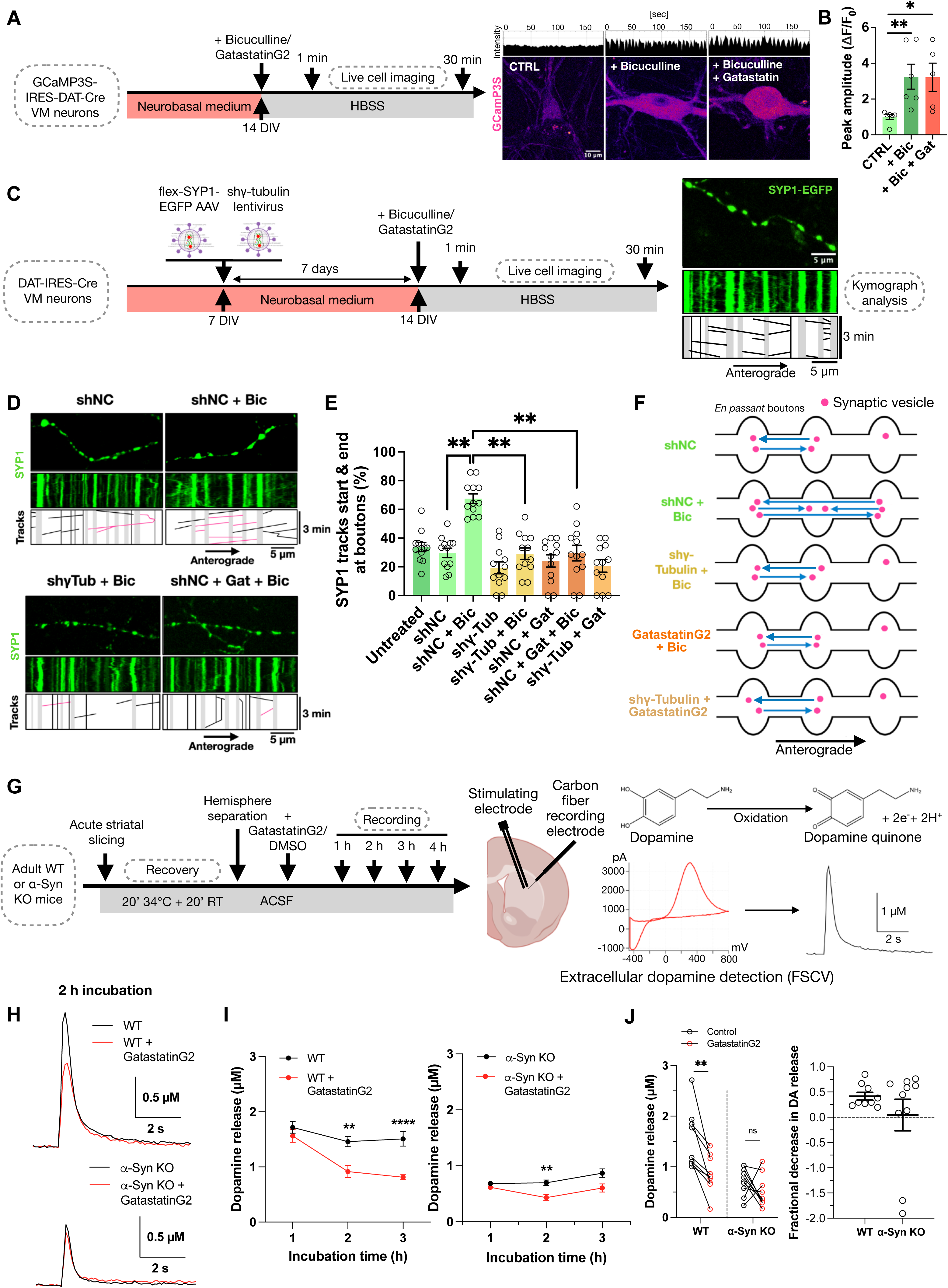
Presynaptic microtubule nucleation promotes interbouton synaptic vesicle exchange and dopamine release in dopaminergic axons. **(A)** Schematic of the experimental design with representative frames from live-imaging and fluorescence intensity profiles of GCamP3S-DAT-IRES-Cre neurons upon bicuculline stimulation with or without the γ-tubulin inhibitor GatastatinG2. **(B)** Peak amplitudes of GCamP3S fluorescence in GCamP3S-DAT-IRES-Cre neurons treated as in (A). Data are mean ± SEM; ** p<0.01, Kruskal-Wallis non-parametric ANOVA. **(C)** Schematic of the experimental design with representative live imaging frames and kymograph from distal axons of DAT-Cre neurons expressing flexed synaptophysin1 (SYP1)-EGFP. All SYP1 tracks in the kymograph are shown in black. **(D)** Representative kymographs from axons of DAT-Cre neurons expressing flexed SYP1-EGFP after 7 days of lentiviral delivery of shRNA targeting γ-tubulin (shγ-Tub) or non-coding control (shNC), or after acute application of GatastatinG2 (Gat). Recordings were performed at baseline or after bicuculline incubation (shNC + Bic, shγ -Tub + Bic, and shNC + Gat + Bic). For each condition, schematic kymographs illustrate SYP1 puncta in motion (black) or “starting & ending” (purple) relative to stable SYP1^+^ boutons (grey). **(E)** Quantification of starting & ending SYP1^+^ puncta relative to stable SYP1^+^ puncta in axons of DAT-Cre neurons treated as in (D). **(F)** Schematic summary of SYP1 dynamics relative to stable SYP1^+^ puncta in axons of DAT-Cre neurons under the conditions shown in (D). **(G)** Schematic of fast-scan cyclic voltammetry (FSCV) experimental design in oblique horizontal acute striatal slices, illustrating the dopamine oxidation reaction, an example of raw current measurements, and the corresponding extracellular dopamine concentration traces over time. **(H)** Representative extracellular dopamine concentration traces recorded after 2 hours of WT and α-synuclein KO (black) alone or after incubation with GatastatinG2 (red). **(I)** Left: WT (N = 6 animals, n = 14 slices in each treatment group); Right: α-synuclein KO (N = 4 animals, n = 13 slices); striatal dopamine release measured in WT mice by FSCV in acute slices over a 3-hour time course in the presence of GatastatinG2. Data are mean ± SEM; ****p<0.0001, Two-Way ANOVA with Bonferroni post hoc multiple comparison test for each time point versus control. **(J)** Effect of GatastatinG2 on evoked DA release in WT and α-syn KO slices following 2–3 h of incubation. Left: each point represents an individual slice; lines connect control and treated slices from the same animal recorded at the same time using the same recording electrode. Right: fractional decrease in DA release, calculated for each matched pair as (control - GatastatinG2)/control; negative values indicate increased release. Each point represents one slice pair; horizontal bars and error bars indicate mean ± SEM. n = 9 WT and 10 α-syn KO slice pairs. ** p < 0.01; ns, not significant; paired t-test for control versus GatastatinG2 within each genotype.

We then tested whether γ-tubulin controls SV trafficking between presynaptic boutons during tonic activity. Using live imaging of distal axons (>100 µm from the soma) expressing fluorescently tagged SYP1, we visualized SVs and *en passant* boutons while suppressing γ-tubulin by shRNA or with GatastatinG2 (**Fig. 3C, D**). At baseline, interbouton SV exchange, consisting of SVs leaving one bouton and arriving at another, was unchanged across conditions (shNC vs shγ-Tub vs shNC + Gat), consistent with a limited contribution of γ-tubulin-dependent processes to basal SV trafficking. We then examined the effects of increased neuronal activity on SV trafficking. We found that synaptic activation with bicuculline doubled the proportion of SYP1 tracks starting and ending at neighboring boutons (up to 70% of all SYP1 tracks) compared to baseline, indicating that enhanced neuronal activity markedly increases interbouton SV exchange. This effect of neuronal activation revealed a strong dependence of SV transport on γ-tubulin: both knockdown and acute inhibition of γ-tubulin markedly decreased activity-evoked interbouton transport to baseline levels (shNC + Bic vs shγ-Tub + Bic vs shNC + Gat + Bic). The combination of γ-tubulin knockdown and GatastatinG2 inhibition produced no further change in SV trafficking, consistent with their effects converging on a common pathway (**Fig. 3E**). Notably, the proportions of anterograde, retrograde, and bidirectional SYP1 tracks did not differ significantly across conditions, with no detectable shift in the directionality of SV transport (**Fig. S3A**). We concluded that in dopaminergic neurons, increased activity drives interbouton SV trafficking, and that γ-tubulin acts as a permissive factor for *de novo* presynaptic MT nucleation required for activity-dependent interbouton SV exchange. However, basal trafficking appears to rely on alternative or γ-tubulin-independent cytoskeletal mechanisms (**Fig. 3F**).

To examine whether γ-tubulin-dependent processes support dopamine release from striatal terminals, we performed FSCV in oblique horizontal striatal slices (10° angle), a preparation that better preserves axonal fibers along the plane of the medial forebrain bundle than conventional coronal sections (Chuhma et al., 2004; Kim et al., 2011) (**Fig. 3G**). We found that in WT slices, GatastatinG2 reduced evoked dopamine release, becoming evident after 2–3 h of incubation (**Fig. 3H, I**). Because α-synuclein regulates presynaptic dopamine release and associates with tubulin, we then examined the effect of GatastatinG2 in α-synuclein KO slices. We observed that the fractional reduction was smaller in KO slices, although a significant effect remained detectable at the 2 h time point (**Fig. 3I**). In paired comparisons within the 2–3 h incubation window, GatastatinG2 significantly reduced release in WT slices, whereas the small reduction in KO slices did not reach statistical significance (**Fig. 3J**). No significant treatment-associated change in dopamine transient half-decay time was detected in either genotype (**Fig. S3B**), indicating that dopamine reuptake activity was not changed under the condition tested. This gradual decline in the WT striatum is consistent with a progressive reduction in functional SV supply, supporting a role for γ-tubulin-dependent presynaptic MT nucleation in SV trafficking and replenishment that may help sustain dopaminergic neurotransmission. These findings prompted us to investigate the functional relationship between this process and α-synuclein.

### α-Synuclein regulates evoked stable dopamine release, activity-dependent presynaptic MT nucleation, and interbouton synaptic vesicle motility

*In vivo* work has revealed activity-dependent effects of α-synuclein on dopamine release, indicating that it dynamically regulates presynaptic function and both long- and short-term plasticity (Shashaank et al., 2023; Somayaji et al., 2020). The apparently attenuated effect of GatastatinG2 in α-synuclein KO striatal slices (**Fig. 3H–J**) raised the possibility that α-synuclein and γ-tubulin-dependent MT nucleation contribute to a shared presynaptic process. To further examine this relationship, we directly compared the drug-free WT and α-synuclein KO recordings from the same FSCV experimental series presented in Fig. 3. We found that stimulated α-synuclein KO slices with single pulse exhibited reduced evoked dopamine release compared with WT controls (**Fig. 4A, B**), indicating a release deficit in striatal terminals, which was still present after a train of 5 stimuli at 40Hz (**Fig. S3C**). The similar dopamine-release phenotypes observed following α-synuclein loss or MT nucleation inhibition, together with the reported association of α-synuclein with tubulin at dopaminergic synapses (Amadeo et al., 2021), suggested a functional link between α-synuclein-dependent presynaptic organization and activity-evoked MT nucleation in dopaminergic axons. To directly test this possibility, we reduced α-synuclein levels using lentiviral shRNA and monitored MT dynamics by live imaging (**Fig. 4C**). α-Synuclein knockdown did not alter MT dynamics at baseline (shNC vs shSNCA) but completely abolished the activity-evoked increase in comet density induced by bicuculline stimulation (shNC + Bic vs shSNCA + Bic), indicating that α-synuclein is required for the activity-dependent increase in MT dynamics (**Fig. 4D, E**). This effect was largely driven by a loss of presynaptic MT initiation, reflected by fewer comets initiating at stable presynaptic puncta (**Fig. 4F**), closely phenocopying γ-tubulin suppression. Importantly, α-synuclein depletion did not detectably alter bouton density, size, or spacing, or the measured γ-tubulin fluorescence intensity and its overlap with synapsin1/2 (**Fig. S4A, C**). The loss of activity-evoked MT initiation therefore occurred without a detectable change in overall bouton organization or γ-tubulin synaptic localization.

**Figure 4.**
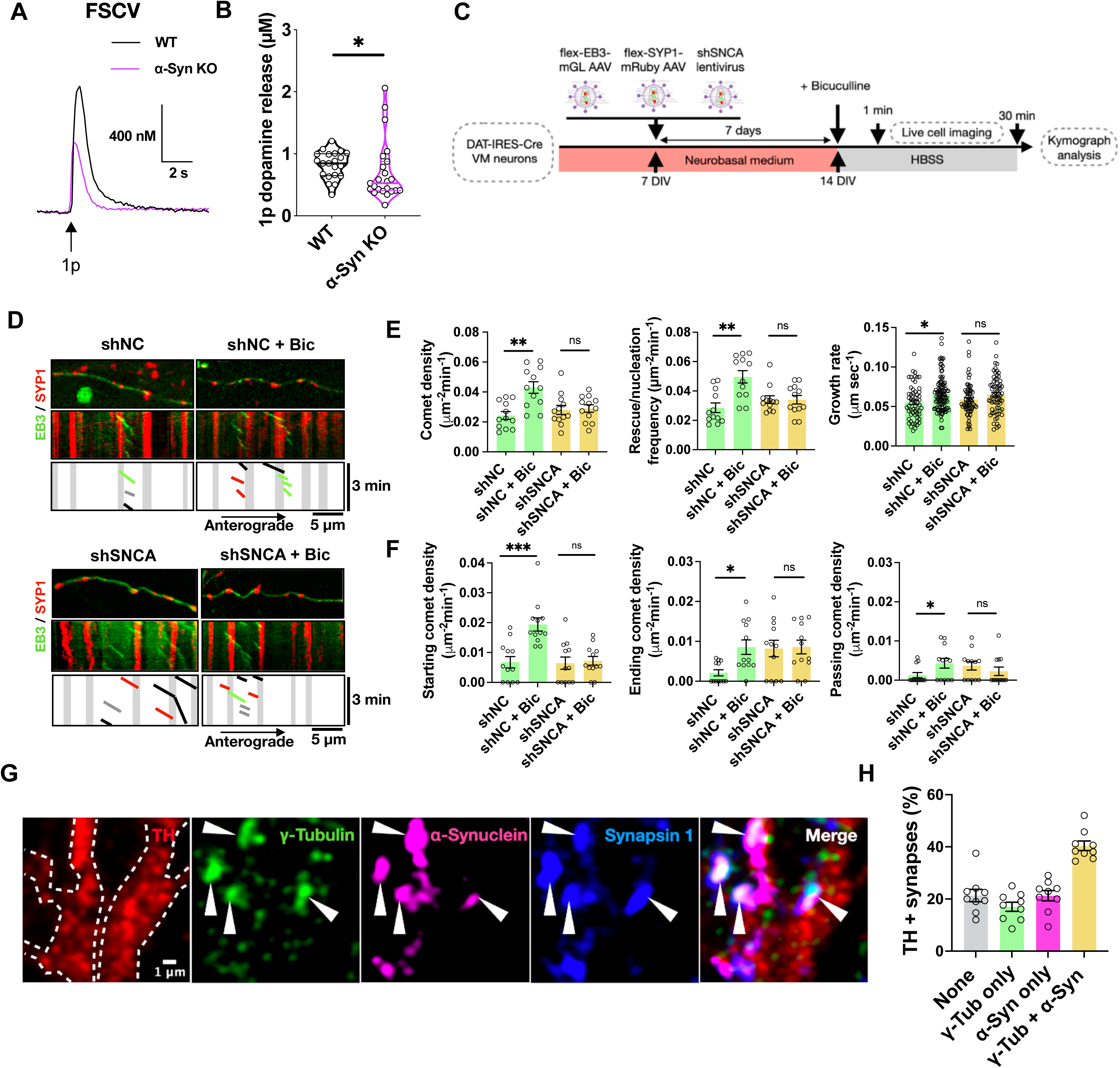
α-Synuclein promotes activity-evoked presynaptic microtubule nucleation at *en passant* boutons in dopaminergic neurons. **A)** Representative FSCV traces of dopamine release evoked by single stimulation and corresponding 1p dopamine release in acute oblique horizontal striatal slices from WT and α-synuclein KO mice. Control traces (black line) are the same as in Fig. 3H. **(B)** Quantification of pooled dopamine release evoked by a single stimulation from WT and α-synuclein KO mice. WT N=6; α-synuclein KO N=4; n= at least 90 recording spots. Data are mean ± SEM; * p<0.01, Mann-Whitney nonparametric test. **(C)** Schematic of the experimental design. **(D)** Representative kymographs from axons of DAT-Cre neurons expressing flexed EB3-mGreenLantern and SYP1-mRuby after 7 days of lentiviral delivery of shRNA targeting α-synuclein (shSNCA) or non-coding control (shNC), at baseline and after bicuculline incubation (shNC + Bic and shSNCA + Bic). **(E)** Quantification of comet density, rescue/nucleation frequency, and growth rate in axons of DAT-Cre neurons expressing flexed EB3-mGreenLantern and SYP1-mRuby in neurons treated as in (D). Data are mean ± SEM; ** p<0.01, Kruskal-Wallis non-parametric ANOVA. **(F)** Quantification of starting, ending, and passing-through comet density relative to stable SYP1^+^ puncta in DAT-Cre neurons treated as in (D). Controls (shNC and shNC + Bic) for starting comets are the same as in Fig. 2G. Data are mean ± SEM; *** p<0.001, Kruskal-Wallis non-parametric ANOVA. **(G)** Representative single-plane Airy-Scan image of endogenous colocalization of γ-tubulin and α-synuclein at dopaminergic synapses, defined by a combined synapsin1 (pan-synaptic marker) and TH (dopaminergic marker) mask. White arrows indicate synapses containing both γ-tubulin and α-synuclein. **(H)** Classification of 307 dopaminergic synapses (TH^+^ and syn1^+^) into 4 categories: neither γ-tubulin nor α-synuclein, γ-tubulin only, α-synuclein only, or both, based on confocal images acquired from the staining shown in (G).

Consistent with a role for MT nucleation in SV trafficking, α-synuclein knockdown also impaired activity-evoked interbouton vesicle transport, selectively reducing SV tracks connecting neighboring boutons, similar to the phenotype observed following γ-tubulin depletion (**Fig. 3D, E, S4D**). Finally, using high-resolution imaging, we observed that in wild-type cultures, ∼40% of TH+ and synapsin1+ axonal boutons contained both α-synuclein and γ-tubulin, whereas ∼20% contained neither (**Fig. 4G, H**), indicating that these proteins coexist at a substantial proportion of dopaminergic presynaptic sites.

Together, these findings support a model in which α-synuclein and γ-tubulin functionally cooperate to regulate activity-dependent presynaptic MT nucleation and interbouton SV exchange, providing a mechanism that may help sustain dopamine neurotransmission during increased neuronal activity.

### α-Synuclein C-terminus binds to the α/β-tubulin heterodimer and is required for activity-dependent presynaptic microtubule nucleation at en passant boutons in dopaminergic neurons

We hypothesized that α-synuclein may directly bind γ-tubulin and/or α/β-tubulin dimers and that these interactions may contribute to presynaptic MT nucleation, as α/β-tubulin dimers are incorporated into nascent MTs assembled from γ-tubulin nucleation templates. To test this, we mapped α-synuclein regions that interact with tubulin using a peptide spot array spanning full-length human α-synuclein (**Fig. 5A**). We identified two N-terminal segments (AA 9–15 and 21–39) and a region at the C-terminal end of the non-amyloid component (NAC) domain (AA 88–102) that bound both α/β-tubulin dimers and γ-tubulin (**Fig. 5B, C**). In contrast, the distal C-terminal segment (AA 126–140) selectively bound α/β-tubulin dimers (**Fig. 5B, C**) and was the only interaction retained following removal of the α/β-tubulin C-terminal tails (**Fig. S5A**), suggesting engagement of the folded tubulin core rather than nonspecific contacts with the disordered, charged C-terminal tails. These interaction domains are summarized schematically in **Fig. 5D**. Notably, the α-synuclein C-terminus also mediates interactions with presynaptic partners involved in SV recycling, including VAMP2, while binding to the synapsin E-domain enables α-synuclein-dependent regulation of SV trafficking (Atias et al., 2019; Stavsky et al., 2024; Sun et al., 2019). Moreover, the tubulin-binding segment identified here contains Ser129, whose activity-dependent phosphorylation promotes interactions with VAMP2 and synapsin that mediate α-synuclein’s physiological regulation of SV recycling (Parra-Rivas et al., 2023; Ramalingam et al., 2023). Together, these observations raised the possibility that this segment also couples neuronal activity to presynaptic MT nucleation.

**Figure 5.**
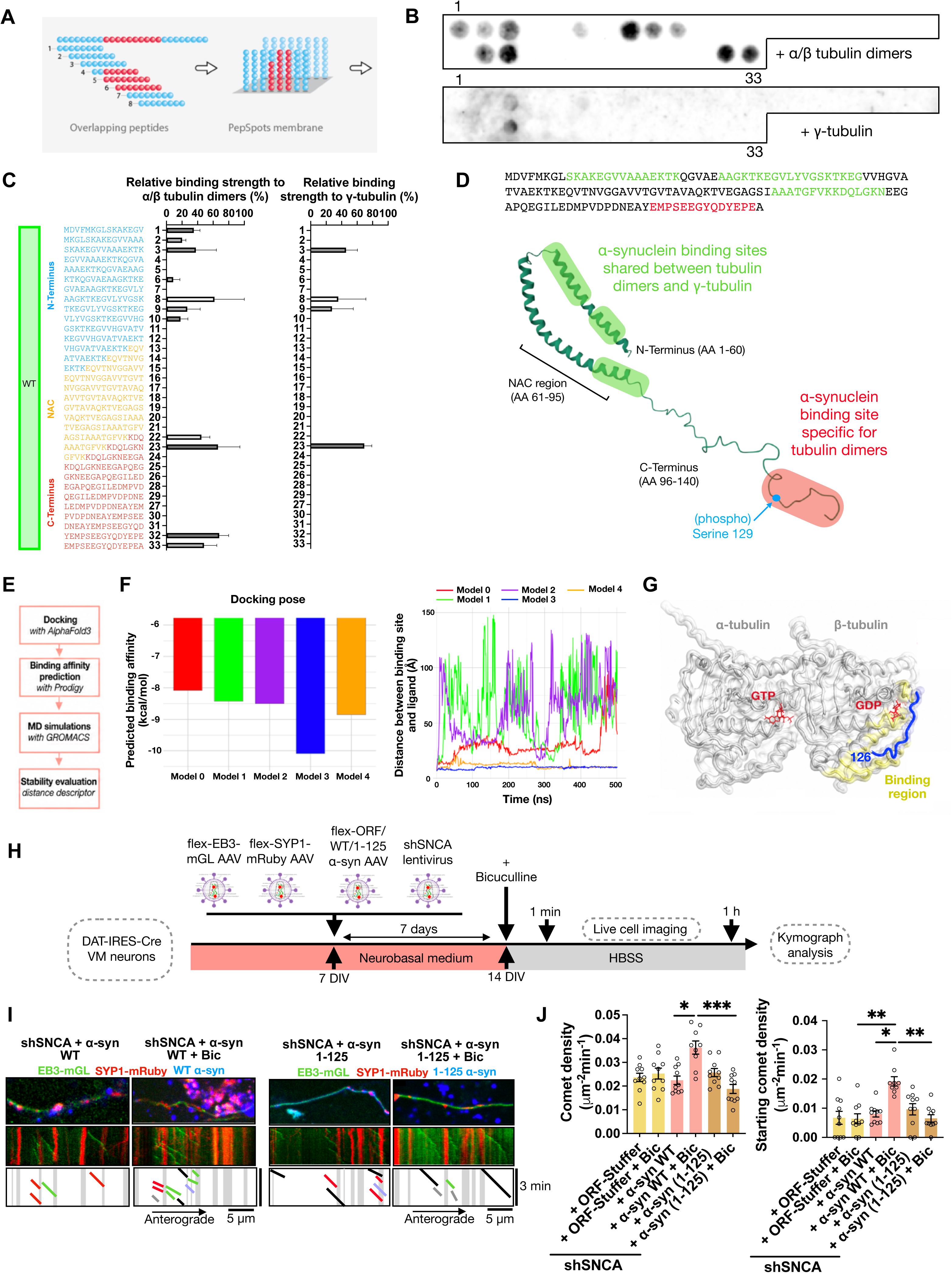
α-Synuclein C-terminus binds to the α/β-tubulin heterodimer and is required for activity-dependent presynaptic microtubule nucleation at en passant boutons in dopaminergic neurons. **(A)** Schematic of the pep-spot array approach, in which WT human α-synuclein is divided into 33 overlapping 15 AA peptides, N-acetylated and immobilized on a cellulose membrane. **(B)** Representative pep-spot results from membranes incubated with either 2.5µg/ml native soluble porcine α/β tubulin dimers or recombinant native human γ-tubulin and developed with infrared-conjugated antibodies. **(C)** Sequence of the 33 WT α-synuclein peptides and corresponding binding strength (%) to soluble α/β tubulin dimers or γ-tubulin. Data are mean ± SEM; n=2 membranes with 2 independent preparations of purified α/β tubulin or γ-tubulin. **(D)** Full-length human WT α-synuclein amino acid sequence with regions that bind both tubulin dimers and γ-tubulin highlighted in green and the tubulin dimer-specific binding region highlighted in red, shown above the 3D lipid-bound structure of human WT α-synuclein (PDB 1XQ8) with the same regions mapped. **(E)** Schematic of the computational pipeline used in this study, including AlphaFold3-based docking, binding affinity estimation, and molecular dynamics simulations. **(F)** Time evolution of the distance between the geometric center of the tubulin binding site (as defined in the initial docking configuration) and the geometric center of the peptide. Distances were calculated every 1 ns throughout the simulations. Each color represents one of the five top-ranked docking poses predicted by AlphaFold3. **(G)** Cartoon representation of the simulated complex, showing the two tubulin chains α/β, α-synuclein peptide 33 (AA 126-140) (model 3), and the bound GDP and GTP ligands. **(H)** Schematic of the experimental design. **(I)** Representative kymographs from axons of DAT-Cre neurons expressing flexed EB3-mGreenLantern and SYP1-mRuby after 7 days of lentiviral delivery of shRNA targeting α-synuclein (shSNCA) at baseline or after bicuculline incubation. α-Synuclein knockdown was rescued with either the control vector (TagBFP2-T2A-ORF-Stuffer) (yellow), WT α-synuclein (pink), or a C-terminally truncated α-synuclein (1-125) (brown). **(J)** Quantification of total and starting comet density relative to stable SYP1^+^ puncta in DAT-Cre neurons treated as in (H). Data are mean ± SEM; *** p<0.001, Kruskal-Wallis non-parametric ANOVA.

To further examine how the α-synuclein C-terminal region interacts with tubulin, we next performed molecular docking of peptide 33 (AA 126–140) onto the α/β-tubulin dimer (**Fig. 5E**). Among five independently predicted docking poses, one configuration displayed a stronger predicted binding affinity (−10.1 kcal/mol) than the others (second-best: −8.85 kcal/mol). Molecular dynamics simulations further that this high-affinity complex remained stably associated with the tubulin surface over time, maintaining a short, relatively constant distance between the peptide and its predicted binding site, whereas the alternative poses showed increased separation and variability (**Fig. 5F**). In this preferred configuration, the C-terminal region of α-synuclein localized at the interface between α- and β-tubulin within the heterodimer (**Fig. 5G**), in close proximity to the exchangeable GTP-binding site of β-tubulin, suggesting a potential structural basis for the reported effects of α-synuclein on tubulin polymerization dynamics *in vitro* (Cartelli et al., 2016).

We then examined whether the α-synuclein C-terminus was functionally required for activity-dependent presynaptic MT nucleation. Following endogenous α-synuclein knockdown (shSNCA), neurons were transduced with either ORF-Stuffer, WT α-synuclein, or a C-terminally truncated variant (α-synuclein 1–125) lacking the putative tubulin-binding segment (**Fig. 5H**). Live imaging and kymograph analyses revealed that the C-terminal truncation did not appreciably alter baseline MT dynamics (ORF-stuffer vs α-syn WT vs α-syn 1–125), but failed to rescue the activity-evoked increase in comet density induced by bicuculline stimulation (ORF-stuffer + Bic vs α-syn WT + Bic vs α-syn 1–125 + Bic) (**Fig. 5I, J**), indicating that the C-terminal residues 126-140 of α-synuclein are essential for activity-dependent presynaptic MT initiation.

### PLK2-dependent α-synuclein Ser129 phosphorylation couples synaptic activity to presynaptic microtubule nucleation at active boutons in dopaminergic neurons

Given the requirement for the α-synuclein C-terminal region in activity-dependent presynaptic MT nucleation, we next examined the role of Ser129 phosphorylation, a dynamic PLK2-dependent modification in the C-terminal region that is induced by neuronal activity and enriched at presynaptic boutons (Parra-Rivas et al., 2023; Ramalingam et al., 2023). To determine whether Ser129 phosphorylation modulates the interaction of α-synuclein with tubulin, we first performed peptide-spot binding assays using peptides spanning the α-synuclein C-terminal 126–140 region in WT, phospho-null (S129A), or phosphomimetic (S129D) configurations (**Fig. 6A**). While binding to γ-tubulin was minimal across all conditions, S129D-containing peptides displayed markedly enhanced binding to α/β-tubulin dimers compared to WT or S129A peptides, particularly within peptides 32–33 (**Fig. 6A, B**). S129A showed tubulin binding comparable to WT, consistent with their shared non-phosphorylated state. These results identify the α-synuclein C-terminal 126–140 region as a phosphorylation-sensitive α/β-tubulin-binding domain, suggesting that phosphorylation at Ser129 enhances this interaction.

**Figure 6.**
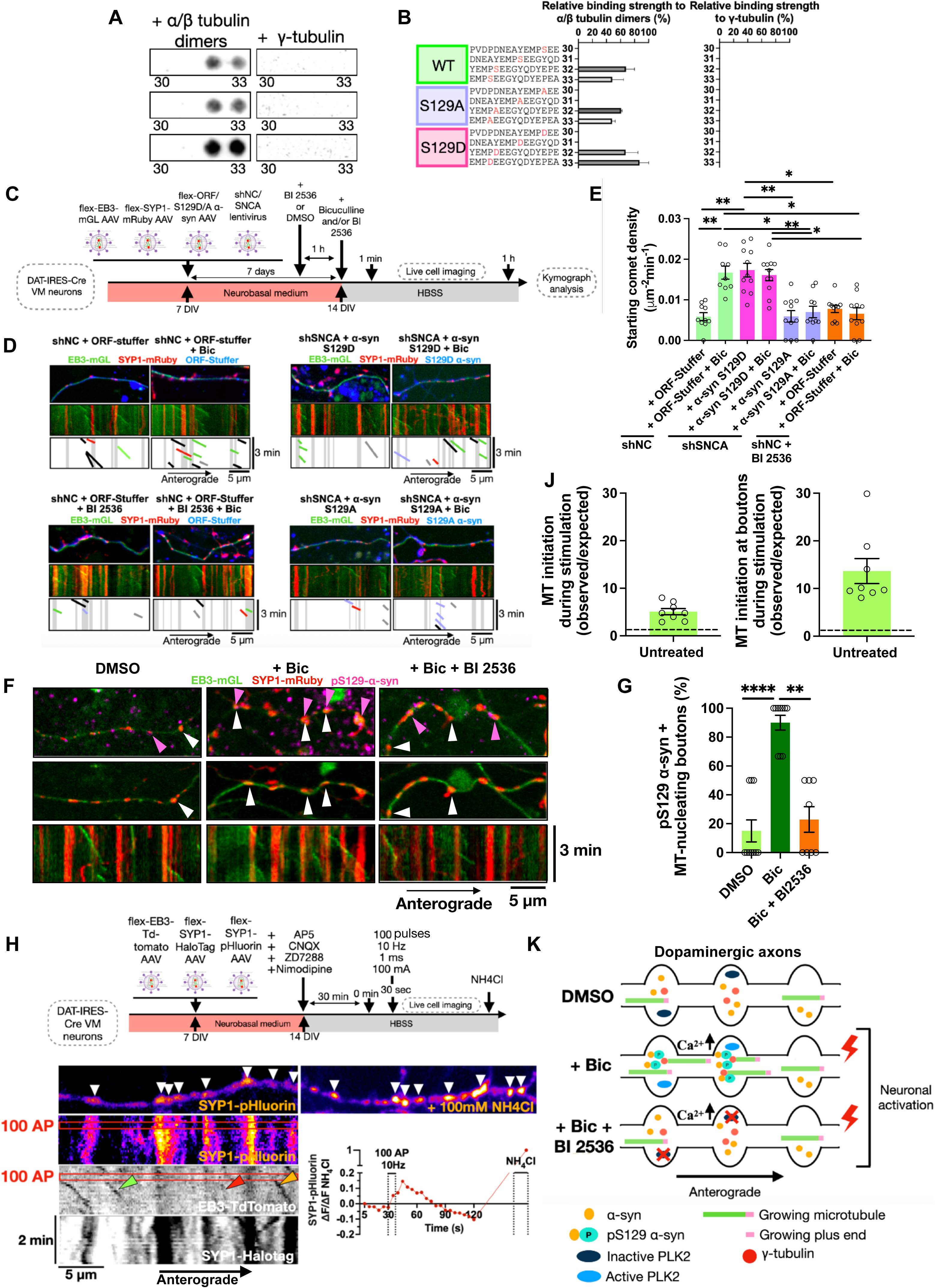
Activity-dependent α-synuclein Ser129 phosphorylation drives presynaptic microtubule initiation at *en passant* boutons in dopaminergic neurons. **(A)** Representative pep spot results for the last four peptides (30-33) of human WT, S129A, and S129D α-synuclein, incubated with 2.5µg/ml native soluble porcine α/β tubulin dimers or recombinant native human γ-tubulin, and developed with infrared-conjugated antibodies. **(B)** Sequence of the last four peptides (30-33) of WT, S129A, and S129D α-synuclein, with the position of S129 (WT), S129A, or S129D highlighted in red, and corresponding binding strength (%) to soluble α/β tubulin dimers or γ-tubulin. Data are mean ± SEM; n=2 membranes with 2 independent preparations of purified α/β tubulin or γ-tubulin. **(C)** Schematic of the experimental design. **(D)** Representative kymographs from axons of DAT-Cre neurons expressing flexed EB3-mGreenLantern and SYP1-mRuby after 7 days of lentiviral delivery of shRNA targeting α-synuclein (shSNCA) or non-coding control (shNC) at baseline or after bicuculline incubation. **(E)** Quantification of starting comet density relative to stable SYP1^+^ puncta in DAT-Cre neurons treated as in (D). The shNC condition expressed the control vector (TagBFP2-T2A-ORF-Stuffer) (green), while α-synuclein knockdown was rescued with phospho-mimetic S129D (magenta) or phospho-null S129A α-synuclein (purple). The control condition was also incubated with the PLK2 inhibitor BI 2536 (orange). Data are mean ± SEM; ** p<0.01, Kruskal-Wallis non-parametric ANOVA. **(F)** Representative kymographs from axons of DAT-Cre neurons expressing flexed EB3-mGreenLantern and SYP1-mRuby at baseline, after bicuculline incubation, or bicuculline + PLK2 inhibitor BI 2536. White arrows indicate the synapses where MT nucleation occurs during the time-lapse movie. Top panel, post-fixation immunostaining with antibody against pS129-α-synuclein and relocation of the imaged axons. Pink arrows indicate boutons positive for pS129-α-synuclein puncta. **(G)** Quantification of MT-nucleating boutons positive for pS129-α-synuclein staining (%) in the imaged DAT-Cre neurons treated as in (F). Data are mean ± SEM; **** p<0.0001, Kruskal-Wallis non-parametric ANOVA. **(H)** Schematic of the experimental design. **(I)** Representative images and kymographs from axons of DAT-Cre neurons expressing flexed EB3-TdTomato, SYP1-pHluorin and SYP1-HaloTag at baseline or after bicuculline incubation. Red boxes on the kymograph indicate the 2 frames (10 sec) in which the electrical stimulation was delivered. The red arrow indicates EB3 comets initiating at a synapse during exocytosis; the bright green arrow indicates a comet initiating during electrical stimulation; and the yellow arrow indicates EB3 comets initiating at a synapse undergoing exocytosis during stimulation with 100AP at 10 Hz. On the right, a representative image after incubation with 100 mM NH4Cl and the corresponding ΔF/ΔF NH4Cl curve for the same axon shown on the left. Each dot in the curve represents a time-lapse movie frame. White arrows indicate each synapse containing SVs, highlighted by bright fluorescence after alkalization with NH_4_Cl. **(J)** Observed/expected ratio (fold change) of comet initiation during 100 AP stimulation along the entire axon (left) or restricted to bouton locations (right). The dashed line indicates Y=1 if the expected and observed values coincide. Each point represents a neurite analyzed. **(K)** Schematic summary of neuronal-activity-driven PLK2-dependent α-synuclein Serine 129 phosphorylation and presynaptic MT nucleation in DAT-Cre neurons.

We next examined the functional relevance of Ser129 phosphorylation in activity-dependent presynaptic MT nucleation using phosphomimetic (S129D) and phospho-null (S129A) α-synuclein mutants in live imaging experiments (**Fig. 6C**). S129D approximates the negative charge of phosphorylation, whereas S129A cannot be phosphorylated at this position, which is targeted by PLK2, a major α-synuclein Ser129 kinase in neurons (Ramalingam et al., 2023). Live imaging and kymograph analyses revealed that S129D elevated MT dynamics at baseline (ORF-stuffer vs α-syn S129D) and did not further enhance starting comet density upon bicuculline stimulation (ORF-stuffer + Bic vs α-syn S129D + Bic), consistent with a constitutively “active” state of presynaptic MT initiation. In contrast, S129A failed to support activity-evoked comet formation (ORF-stuffer + Bic vs α-syn S129A + Bic), indicating that the ability to phosphorylate Ser129 is required for activity-dependent presynaptic MT initiation (**Fig. 6D, E**).

To determine whether these effects depended on endogenous PLK2-mediated Ser129 phosphorylation, we next applied the PLK2 inhibitor BI2536 before and during bicuculline stimulation (**Fig. 6C**). PLK2 inhibition phenocopied the phospho-null S129A mutant, completely abolishing the activity-dependent increase in MT initiation at presynaptic boutons (ORF-Stuffer + Bic vs ORF-Stuffer + BI2536 + Bic) (**Fig. 6D, E**), supporting a role for PLK2-dependent α-synuclein phosphorylation in coupling neuronal activity to presynaptic MT initiation.

We then asked whether boutons exhibiting activity-dependent MT initiation were selectively enriched in pS129-α-synuclein. Post-imaging fixation, pS129-α-syn immunostaining, and quantitative analyses were performed on the same live-imaged axonal segments and presynaptic boutons used for MT intiation measurements, enabling the bouton-by-bouton comparison of MT initiation behavior and pS129-α-syn enrichment (**Fig. 6F**). We found that bicuculline increased pS129-α-syn levels at boutons, consistent with reports linking pS129-α-syn to activity-dependent presynaptic function (Parra-Rivas et al., 2023; Ramalingam et al., 2023) (**Fig. S6B**), and increased the fraction of MT-initiating boutons positive for pS129-α-synuclein, whereas BI2536 suppressed both effects (**Fig. 6G**).

In addition, proximity ligation assay (PLA) revealed a close proximity between endogenous α-synuclein and γ-tubulin at dopaminergic synapses. Bicuculline increased the density of synaptic pS129-α-synuclein/γ-tubulin PLA puncta, whereas total α-synuclein/γ-tubulin PLA puncta density did not change significantly (**Fig. S6D**). To examine the temporal relationship between MT initiation and synaptic exocytosis, we simultaneously imaged EB3-TdTomato, SYP1-pHluorin (reporting SV exocytosis), and SYP1-HaloTag (labeling presynaptic boutons) in dopaminergic axons during electrical stimulation (**Fig. 6H, I**). Stimulation elicited SV fusion accompanied by MT initiation events at boutons. Importantly, MT initiation occurred preferentially during the stimulation window, exceeding the frequency expected from the duration of stimulation alone. This enrichment persisted when bouton location was incorporated into the analysis (**Fig. 6J**). These measurements reveal a previously unrecognized temporal association between activity-evoked SV exocytosis and local presynaptic MT initiation, complementing the association between MT initiation and pS129-α-synuclein observed by post-imaging immunostaining. Together, these findings indicate that neuronal activity promotes PLK2-dependent α-synuclein S129 phosphorylation and its preferential enrichment at MT-initiating boutons. They identify S129 as a critical regulatory switch for presynaptic MT nucleation and reveal that MT initiation is temporally coupled to synaptic activity at sites of SV release in dopaminergic axons (**Fig. 6K**).

### PD mutants A53T, A30P, and E46K α-synuclein differentially disrupt presynaptic microtubule initiation and growth at en passant boutons in dopaminergic neurons

Having identified α-synuclein as a regulator of activity-evoked presynaptic MT initiation, we asked whether the PD-associated variants A53T, A30P, and E46K alter this function. Peptide-array mapping revealed mutation-specific changes in tubulin binding. A30P and E46K exhibited expanded binding patterns to soluble α/β-tubulin and γ-tubulin across mutation-containing peptides relative to the corresponding WT sequences, with E46K producing a particularly pronounced gain in binding, whereas peptides spanning A53T showed little or no detectable binding in either configuration (**Fig. 7A–C**). Thus, these mutations exert distinct effects on the recognition of α/β-tubulin and γ-tubulin by mutation-containing peptides.

**Figure 7.**
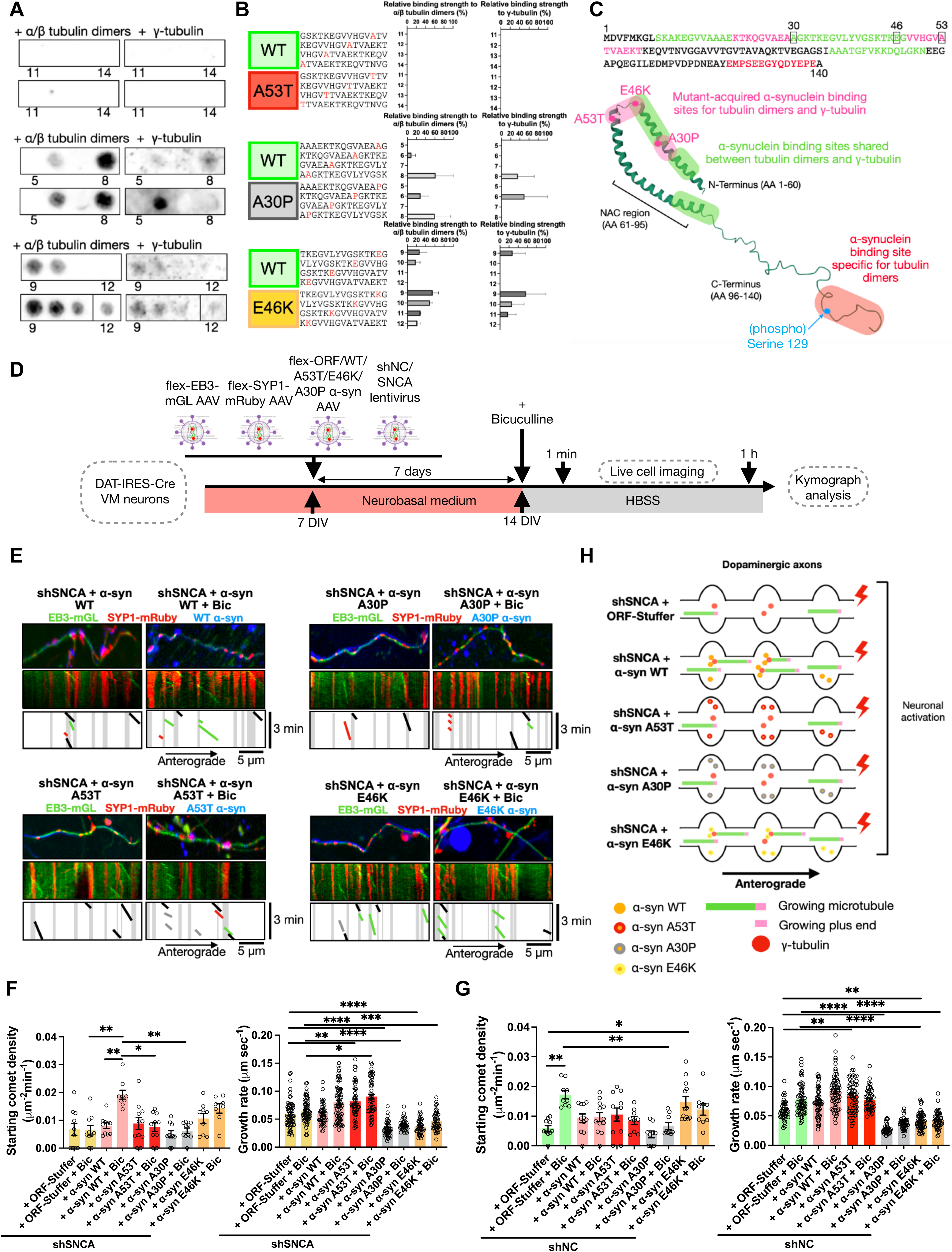
PD mutants A53T, A30P, and E46K α-synuclein differentially disrupt presynaptic microtubule initiation and growth at *en passant* boutons in dopaminergic neurons. **(A)** Representative pep-spot results from membrane mapping of the PD-associated α-synuclein mutants A53T, A30P, and E46K with four peptides each, covering the 12 aa region before and after the mutation on the α-synuclein sequence. Membranes were incubated with either 2.5 µg/ml native soluble porcine α/β tubulin dimers or recombinant native human γ-tubulin and then developed with infrared-conjugated antibodies. **(B)** Sequence of the four peptides covering each PD-associated α-synuclein mutant, with the position of A53T, A30P, and E46K highlighted in red, and corresponding binding strength (%) to soluble α/β tubulin dimers or γ-tubulin. Data are mean ± SEM; n=2 membranes with 2 independent preparations of purified α/β tubulin or γ-tubulin. **(C)** Full-length human WT α-synuclein amino acid sequence with regions that bind both tubulin dimers and γ-tubulin highlighted in green, the tubulin dimer-specific binding region highlighted in red, and the tubulin/γ-tubulin acquired binding region of PD mutants highlighted in pink, shown above the 3D lipid-bound structure of human WT α-synuclein (PDB 1XQ8) with the same regions mapped. **(D)** Schematic of the experimental design. **(E)** Representative kymographs from axons of DAT-Cre neurons expressing flexed EB3-mGreenLantern and SYP1-mRuby after 7 days of lentiviral delivery of shRNA targeting α-synuclein (shSNCA) at baseline or after bicuculline incubation. α-Synuclein knockdown was rescued with either control vector (ORF-Stuffer), WT (pink), or PD mutant α-synuclein A53T (red), A30P (grey), or E46K (yellow). **(F)** Quantification of starting comet density relative to stable SYP1^+^ puncta and total growth rate in DAT-Cre neurons treated as in (D) after rescue of α-synuclein knockdown (shSNCA). ORF-Stuffer and WT rescue values are the same as in Fig. 5J. Data are mean ± SEM; **** p<0.0001, Kruskal-Wallis non-parametric ANOVA. **(G)** Quantification of starting comet density relative to stable SYP1^+^ puncta and total growth rate in DAT-Cre neurons treated as in (D) after overexpression of α-synuclein WT and PD mutants (shNC). ORF-Stuffer control values are the same as in Fig. 6E. Data are mean ± SEM; **** p<0.0001, Kruskal-Wallis non-parametric ANOVA. **(H)** Schematic summary of the loss of function of α-synuclein PD mutants on activity-dependent presynaptic MT nucleation in DAT-Cre neurons.

We next expressed the variants following knockdown of endogenous α-synuclein and measured presynaptic MT initiation before and after bicuculline stimulation (**Fig. 7D–F**). Whereas WT α-synuclein restored the activity-evoked increase in starting comet density, A53T and A30P failed to restore this response. E46K showed a nonsignificant trend toward higher baseline comet density, with a smaller additional increase upon stimulation (**Fig. 7F**). To determine whether these variants also alter MT initiation in the presence of endogenous α-synuclein, we next examined their effects under overexpression conditions. Expression of A53T and A30P in the presence of endogenous α-synuclein also blunted the activity-associated increase in presynaptic MT initiation. In contrast, E46K overexpression significantly increased basal initiation to levels approaching those observed following stimulation (**Fig. 7G**), consistent with a partially constitutive effect on MT initiation. This elevation coincided with enhanced tubulin binding to E46K-containing peptides, suggesting that altered tubulin interactions may contribute to increased initiation under basal conditions. E46K therefore disrupted the normal activity dependence of presynaptic MT initiation through a mechanism that is distinct from the deficient evoked response observed with A53T and A30P. Interestingly, WT overexpression also attenuated activity-evoked presynaptic MT initiation (**Fig. 7G**), suggesting that this process is sensitive not only to α-synuclein mutations but also to increased protein abundance, a perturbation relevant to familial PD caused by SNCA gene duplication or triplication (Chartier-Harlin et al., 2004; Singleton et al., 2003).

Beyond their effects on initiation, A30P and E46K markedly slowed MT growth in both rescue and overexpression conditions (**Fig. 7F, G**). This additional phenotype coincided with their higher binding to tubulin, whereas A53T showed neither a comparable increase in binding nor a comparable slowing of growth relative to WT α-synuclein (**Fig. 7A, B, F, G**). One possible explanation is that the additional interactions detected in A30P- and E46K-containing peptides favor tubulin retention in complexes that are less readily incorporated into growing MTs, although neither sequestration nor reduced dimer availability was directly measured. Together, these findings reveal that disruption of the normal activity-dependent control of presynaptic MT initiation is a consequence of PD-associated α-synuclein mutations, despite their distinct effects on MT dynamics. Whereas A53T and A30P impair activity-evoked initiation, E46K increases basal initiation, suggesting a loss of temporal coupling between neuronal activity and MT assembly. The additional growth defects associated with A30P and E46K further indicate that these mutations differentially affect MT initiation and elongation (**Fig. 7H**).

## DISCUSSION

α-Synuclein expression has long been known to regulate the kinetics of evoked dopamine release (Abeliovich et al., 2000; Oliveira et al., 2021; Shashaank et al., 2023; Somayaji et al., 2020; Sulzer & Edwards, 2019), but how it does so remains unknown. Here, we uncover a previously unrecognized physiological function of α-synuclein in coupling neuronal activity to local MT nucleation at dopaminergic presynaptic boutons. We find that MT initiation in dopaminergic axons preferentially occurs at sites of evoked SV release, revealing a spatial and temporal relationship between presynaptic cytoskeletal remodeling and neurotransmission. We further identify PLK2-dependent phosphorylation of α-synuclein at S129 as a molecular switch that promotes γ-tubulin-dependent MT nucleation, supporting interbouton SV redistribution and regulating dopamine release. Together, these findings suggest a novel regulatory mechanism through which α-synuclein coordinates local SV trafficking with the demands of synaptic activity.

Similar to our prior report in excitatory hippocampal neurons (Qu et al., 2019), we find that dopaminergic axons maintain canonical plus-end-out MT polarity, and that neuronal activity boosts overall dynamic MT density primarily through *de novo* nucleation at γ-tubulin in presynaptic boutons. We find that in dopamine axons, acute γ-tubulin depletion leaves baseline comet density largely unchanged but abolishes the activity-dependent rise in initiating comets at boutons, supporting a model in which γ-tubulin-dependent nucleation is the dominant mechanism for activity-evoked MT initiation at mature dopaminergic synapses. Importantly, simultaneous imaging of MT dynamics and SV exocytosis reveals that MT initiation is enhanced during periods of evoked SV fusion, suggesting that presynaptic MT nucleation is not simply a constitutive mechanism for maintaining axonal architecture but an activity-regulated process linked to local synaptic demand. These results indicate that at dopaminergic synapses, activity-evoked nucleation of dynamic MTs at boutons correlates with increased SV tracks that start and end at neighboring boutons, and are consistent with a role for newly formed MT segments as on-demand tracks for local vesicle exchange that facilitate the replenishment of active release sites during periods of increased synaptic demand. Our findings align with emerging roles of newly formed dynamic MTs in the development and maintenance of presynaptic sites in several models (Aiken & Holzbaur, 2024; Park et al., 2023; Xie et al., 2026). Such spatially restricted cytoskeletal remodeling may be particularly important in dopaminergic neurons, which have extensive axonal arborization and maintain both active and silent presynaptic sites that may impose distinctive demands on SV distribution.

Our data identify α-synuclein and its phosphorylation at Serine 129 as key regulators of presynaptic MT nucleation in dopaminergic axons. α-Synuclein knockdown abolishes activity-induced presynaptic MT nucleation and produces a vesicle transport phenotype closely resembling γ-tubulin depletion, with a selective loss of starting and ending tracks at dopaminergic boutons but preserved directionality. These convergent phenotypes support the participation of the two proteins in a shared presynaptic process. The smaller depression in dopamine release following GatastatinG2 treatment in α-synuclein KO slices is compatible with this functional convergence, although residual drug sensitivity remained detectable, and because dopamine release was already reduced in KO slices, a restricted dynamic range may also contribute to this pattern. Together with our biochemical evidence for α-synuclein binding to γ-tubulin, these findings support a model in which α-synuclein and γ-tubulin functionally cooperate to regulate activity-dependent presynaptic MT nucleation. α-Synuclein also interacts with the SV membrane protein synaptobrevin-2/VAMP2 to promote vesicle clustering and regulate recycling (Diao et al., 2013; Sun et al., 2019), and with synapsin III to control SV pool organization in dopaminergic neurons (Atias et al., 2019; Stavsky et al., 2024; Zaltieri et al., 2015). Together with its interactions with γ-tubulin and soluble α/β-tubulin, these vesicle-associated functions raise the possibility that α-synuclein coordinates SV organization with local MT assembly in response to neuronal activity. By coupling vesicle availability with the formation of MT tracks, α-synuclein may facilitate SV redistribution between neighboring boutons, linking cytoskeletal remodeling to presynaptic platicity. Whether this coordination requires simultaneous or sequential engagement of its vesicular and tubulin partners remains an important question for future investigation.

Interestingly, α-synuclein and γ-tubulin colocalize in many dopaminergic presynaptic boutons, while other boutons lack one or both proteins, indicating a molecular heterogeneity in presynaptic sites that may reflect differences in release competence or active zone composition. This feature may be associated with active and silent synapses in dopaminergic axons (C. Liu et al., 2018; Pereira et al., 2016). Whether the molecular composition of individual boutons determines their competence for activity-dependent MT nucleation, and how this relates to functional release properties, is an important question for future studies.

Our results provide insight into how α-synuclein cooperates with the MT nucleation machinery. Peptide-array mapping identifies N-terminal segments (aa 9–15, 21–39) and the distal NAC region (aa 88–102) that bind both α/β-tubulin dimers and γ-tubulin, while the distal C-terminus (aa 126–140), which includes residue S129, binds tubulin dimers selectively. This interaction is retained after deletion of tubulin C-terminal tails, and so is consistent with recognition of the folded tubulin core. Functionally, re-expression of a C-terminally truncated α-synuclein (1-125) fails to rescue activity-dependent presynaptic MT nucleation after knockdown, indicating that the last 15 residues of α-synuclein are required, in agreement with prior work assigning key synaptic protein interaction roles to the C-terminus (Parra-Rivas et al., 2023; Ramalingam et al., 2023; Stavsky et al., 2024; Sun et al., 2019; Vargas et al., 2025). Consistent with a regulatory role for this region, phosphomimetic S129D enhances α/β-tubulin binding in peptide arrays, whereas the phospho-null S129A mutant fails to support activity-evoked MT initiation (**Fig. 6E**). Together with the enrichment of pS129-α-synuclein at MT-initiating boutons (**Fig. 6F, G**), these findings strongly indicate that activity-dependent S129 phosphorylation promotes tubulin engagement at presynaptic nucleation sites.

These observations further suggest a mechanism in which distinct regions of α-synuclein coordinate interactions with γ-tubulin and soluble α/β-tubulin dimers during the early steps of MT assembly. In this model, the distal C-terminus provides an essential, phosphorylation-sensitive interaction with α/β-tubulin, potentially facilitating its incorporation into newly nucleated MTs. Although the formation of such a complex has not been directly demonstrated, this model extends the previously proposed MT-dynamase function of α-synuclein to an activity-regulated presynaptic mechanism.

γ-Synuclein has been previously described as having MAP-like features, and it may be that tubulin recognition involves both conserved and divergent elements within the synuclein family. The N-terminal α-synuclein segments identified in our peptide arrays, residues 9–15 and 21–39, are highly conserved in γ-synuclein, which associates directly with MTs and promotes tubulin polymerization *in vitro* (Zhang et al., 2011). Both proteins also directly bind to the neuronal-enriched βIII-tubulin, supporting a shared capacity for tubulin interaction, although the binding determinants in γ-synuclein remain unmapped (Toba et al., 2017). In contrast, the distal α-synuclein C-terminal region encompassing residues 126–140 and S129 is poorly conserved in γ-synuclein. In our experiments, deleting the α-synuclein C-terminal residues 126–140 impaired activity-evoked presynaptic MT nucleation without reducing the MT growth rate, supporting a potentially distinct contribution of this region to activity-dependent initiation. Conserved, helix-forming regions may also regulate MT growth, although a functional separation between these tubulin-binding domains remains to be established. Thus, the poorly conserved C-terminal region of α-synuclein may confer a distinct capacity to regulate MT nucleation in response to neuronal activity.

An important implication of our findings is that S129 phosphorylation provides a molecular mechanism linking neuronal activity to presynaptic MT assembly. pS129 α-synuclein is abundant in pathologic inclusions (Spillantini et al., 1997) and has recently been reported as a physiological, activity-dependent signal that boosts α-synuclein’s synaptic protein interactions and enhances synaptic transmission and plasticity *in vivo* (Parra-Rivas et al., 2023; Ramalingam et al., 2023). Consistent with those reports, we find that WT α-synuclein restores activity-evoked MT nucleation, whereas the phospho-mimetic S129D mutant drives elevated baseline nucleation and fails to respond further to stimulation, consistent with a constitutive “on” state. Conversely, rescue with the phospho-null S129A mutant fails to support activity-evoked nucleation. Consistent with this, pharmacological inhibition of PLK2 in neurons expressing endogenous levels of α-synuclein also fails to support activity-evoked nucleation and phenocopies α-synuclein loss. Unlike the shorter MTs reported with S129A *in vitro* (Toba et al., 2017), S129A impaired activity-evoked MT initiation without reducing EB3 comet growth length in our neurons, whereas PLK2 inhibition affected both. This difference may reflect compensation by other neuronal MT regulators or broader effects of kinase inhibition beyond α-synuclein Ser129 phosphorylation. Importantly, post-imaging fixation and immunostaining revealed that neuronal activity increases the fraction of MT-nucleating boutons positive for pS129-α-synuclein, whereas PLK2 activity inhibition suppresses these effects. Neuronal activation also increased synaptic pS129-α-synuclein/γ-tubulin PLA puncta without a detectable change in total α-synuclein/γ-tubulin PLA puncta density, consistent with increased representation of phosphorylated α-synuclein near γ-tubulin at presynaptic sites. Together with the enhanced α/β-tubulin binding observed for S129D-containing peptides, these findings suggest that phosphorylation regulates the ability of α-synuclein to engage soluble tubulin at sites of presynaptic MT nucleation. Thus, neuronal activity does not simply increase pS129-α-synuclein levels globally but promotes its enrichment at boutons undergoing MT initiation. Furthermore, simultaneous imaging of MT dynamics and SV release responses revealed that MT initiation is enriched during the stimulation window accompanying evoked SV exocytosis. Collectively, these data support a model in which neuronal activity engages a PLK2/pS129-α-synuclein pathway that promotes local presynaptic MT nucleation at active synapses. This mechanism provides a molecular link between activity-dependent α-synuclein phosphorylation and local cytoskeletal remodeling, potentially enabling presynaptic boutons to adapt SV trafficking to changing demands for neurotransmitter release.

Our findings suggest that disruption of the physiological control of presynaptic MT nucleation contributes to α-synuclein-associated synaptic dysfunction, and that this occurs with the PD-associated α-synuclein mutations. The A53T and A30P variants compromise the ability of dopaminergic boutons to translate increased neuronal activity into local MT assembly, potentially limiting the redistribution of SVs when synaptic demand rises. A53T retains most of its tubulin-induced folding capability (Cartelli et al., 2016), but shows impaired activity-evoked MT initiation without comparable slowing, suggesting that disrupting the coupling between neuronal activity and local initiation can occur without a general reduction in MT elongation. The E46K mutant displayed a distinct phenotype, significantly increasing basal presynaptic MT initiation in overexpression conditions. The region producing enhanced tubulin binding in E46K overlaps with the 43–63 segment aligned to a tubulin-binding region of stathmin-family proteins (Cartelli et al., 2016), suggesting that this sequence may contribute to tubulin recognition. This enhanced binding may lower the initiation threshold, promoting MT assembly outside periods of increased synaptic demand and weaken the coupling between MT initiation and neuronal activity. In addition, additional tubulin-binding sites detected in A30P- and E46K-containing peptides might favor interactions with soluble tubulin dimers before their incorporation into MTs, potentially sequestering dimers in less assembly-competent complexes and contributing to slower growth.

The impaired activity-dependent response produced by overexpression of WT α-synuclein suggests that this mechanism is vulnerable not only to changes in α-synuclein sequence but also to protein abundance, providing a potential functional connection to familial PD caused by SNCA duplication or triplication: excess α-synuclein might disrupt the balance of interactions required for productive MT initiation, allowing increased abundance and disease-associated mutations to converge on dysregulated activity-dependent presynaptic remodeling.

Our results from α-synuclein mutants indicate that increased tubulin engagement does not necessarily simply produce more effective MT assembly; the functional outcome may depend on where and how these interactions are regulated. Rather than simply impairing MT assembly, these variants appear to disrupt its normal temporal regulation by neuronal activity, suggesting that the appropriate timing of local MT nucleation may be critical for regulating presynaptic function.

Several questions remain regarding the precise relationship between presynaptic MT nucleation and dopamine release. Although our findings support a role for newly nucleated MTs in interbouton SV redistribution, direct visualization of SV transport along these newly formed tracks will be necessary to establish this mechanism. Similarly, it remains to be determined whether S129-dependent MT nucleation directly contributes to the regulation of dopamine release in intact dopaminergic circuits. These questions will help establish how local MT assembly influences synaptic function under physiological conditions and whether its disruption contributes to early dopaminergic dysfunction in PD.

In summary, our findings reveal a physiological function of α-synuclein in coordinating activity-dependent cytoskeletal remodeling with presynaptic neurotransmission in dopaminergic axons. By promoting γ-tubulin-dependent MT nucleation at presynaptic boutons, α-synuclein may enable dopaminergic neurons to dynamically redistribute SVs in response to changing synaptic demands. Disruption of this process by PD-associated α-synuclein mutations suggests that defective presynaptic MT regulation contributes to early synaptic vulnerability in Parkinson’s disease and may represent a novel therapeutic target in PD and related synucleinopathies.

## METHODS

### Experimental models

All protocols and procedures for rats were approved by the Committee on the Ethics of Animal Experiments of Columbia University and by the Guide for the Care and Use of Laboratory Animals of the National Institutes of Health. Time-pregnant Sprague-Dawley rats (Embryonic Day 18) were purchased from Charles River Laboratories for primary hippocampal neuronal cultures. All protocols and procedures for mice followed the National Institutes of Health guidelines and were approved by the Institutional Animal Care and Use Committee of Columbia University and New York State Psychiatric Institute.

#### Primary dopaminergic neuronal cultures

Primary dopaminergic neuronal cultures were prepared as previously described (Staal et al., 2007). Ventral midbrain regions were dissected from newborn mouse pups (DAT-Cre and C57/Bl6J wildtype, Jackson Labs). The tissue was digested with papain (Worthington Biochemical), then gently triturated, spun down, and resuspended in neuronal media. Cells were plated over the glial cells’ monolayer on glass-bottom culture dishes for live imaging and biochemistry assays at a density of 8×^5^ cells/dish. The glial cells were prepared from cortices of newborn-2-day-old Sprague-Dawley rat pups at least 1 week prior to neuronal cultures.

### Method details

#### Lentiviral shRNA silencing

Production of lentiviral particles was conducted using the 2^nd^ generation packaging system as previously described (J. Liu et al., 2014). Briefly, HEK293T cells were co-transfected with lentiviral shRNA constructs and the packaging vectors pLP1, pLP2, and pLP-VSV-G (Invitrogen) using calcium phosphate. 48 h and 72 h after transfection, the virus was collected, filtered through a 0.22 µm filter, and 1 volume of cold Lentivirus Precipitation Solution (Alstem) to every 4 volumes of lentivirus-containing supernatant was added, and the mixture was incubated for at least 6 h at 4°C. The lentivirus-containing supernatant was centrifuged for 30 mins at 3000 rpm and the pellet was resuspended in the minimum volume of neuronal medium recommended by the producer. The concentrated virus was aliquoted and stored at −80°C. Lentiviral construct to knockdown mouse α-synuclein in dopaminergic neurons was purchased from Sigma Aldrich (TRCN0000366591) with the following target sequence GATCCTGGCAGTGAGGCTTAT within a pLKO.1 lentivector. Lentiviral construct to knockdown mouse γ-tubulin in dopaminergic neurons was purchased from Sigma Aldrich (TRCN0000089907) with the following target sequence GCAGCAGCTGATTGACGAGTA within a pLKO.1 lentivector. The vector pLV with a mammalian scramble non-coding sequence was used as a control. The pLKO.1 vector with a noncoding (NC) sequence was used as a control.

#### AAV expression

The control was designed using an ORF-Stuffer (amino acids 2-83 of *E. coli* β-galactosidase) inserted into the mTagBFP2-T2A-ORF-Stuffer construct. All vectors were custom-generated and purchased from Vectorbuilder (maps available upon request). To achieve selective expression only in dopaminergic Cre-positive neurons, all these constructs were designed to be flexed, and a human Synapsin1 promoter controlled the expression of all constructs. All AAVs for live-imaging experiments were used to infect dopaminergic neurons at 7 DIV, and live imaging was performed at 14 DIV.

#### Live imaging of MT and SV dynamics

Lentiviral delivery of shRNA targeting α-synuclein or shNC non-coding control was performed in DAT-Cre ventral midbrain neurons at 7 DIV. On the same day, neurons were co-infected with two AAVs expressing flexed EB3-mGreenLantern and SYP1-mRuby. To rescue α-synuclein knockdown, neurons were co-infected at 7 DIV with flexed AAV expressing mTagBFP2-T2A-α-synuclein (WT, 1-125 C-truncated, S129D, S129A). Control shNC-infected neurons were transduced with a control FLEX AAV (mTagBFP2-T2A-ORFStuffer). The same condition was used to assess the effect of PLK2 inhibition on MT dynamics, with BI 2536 100 nM applied prior to and during bicuculline imaging. Similarly, to measure SV interbouton transport, lentiviral delivery of shRNA targeting γ-tubulin was performed in DAT-Cre ventral midbrain neurons, and on the same day, neurons were co-infected with AAV expressing flexed SYP1-EGFP.

Live cell imaging was performed at 14 DIV, 7 days after AAV infection in complete HBSS media using a 60x oil objective on a Ti2 Eclipse Nikon microscope equipped with dual camera Hamamatsu ORCA-fusion BT C15400, CSU-W1 Yokogawa spinning disk module, and Tokai-Hit live imaging incubator and processed by Nikon NIS Elements software at 2 s/frame for 3 min. Confocal live imaging was performed by acquiring 3 stacks of 0.5 µm each, and the maximum intensity projections of the movies were analyzed using ImageJ. Only stable or wiggling SYP1 puncta with 0.2<x<2.5 µm^2^ size and round or oval shape were considered as boutons. Axons were selected during live imaging based on morphology, exclusive anterograde movement of EB3 comets, and a synaptic-like pattern of SYP1 puncta.

To induce neuronal activity, neurons were washed 1x with complete HBSS, and 20 µM bicuculline or DMSO control was added to the complete HBSS after washes. Movies were acquired 1 min upon treatment. To pharmacologically suppress MT nucleation, neurons were washed 1x with complete HBSS, and 10 µM GatastatinG2 or DMSO control was added to the complete HBSS after washes. Movies were taken 1 min after treatment, up to 30 min. Movies were analyzed in ImageJ. Kymographs were generated by drawing a region in the distal axons (more than 100 µm from the cell body) based on morphology and anterograde movement of EB3-labeled comets. Presynaptic MTs were classified based on their plus-end contacts with stable SYP1-labeled boutons in dopaminergic neurons. In our measurements of EB3 tracks starting or ending at boutons, we also included those that start or end at a bouton and pass through the next distal or proximal boutons. Parameters describing MT dynamics were defined as follows: rescue/nucleation frequency: number of rescue or nucleation events per µm^2^ per min; catastrophe frequency: number of full tracks/total duration of growth; comet density: number of comets per µm^2^ per min; growth length: comet movement length in µm; comet lifetime: duration of growth; growth rate: growth length/comet lifetime. Parameters describing organelle and SV dynamics were defined as follows: % of moving puncta: number of moving puncta/number of total puncta x 100; % of tracks start/end at boutons: number of tracks start or end at boutons/number of total moving tracks x 100; % of tracks start and end at boutons: number of tracks start and end at boutons/number of total moving tracks x 100; % of anterograde tracks: number of anterograde tracks/number of total moving tracks x 100%; of retrograde tracks: number of retrograde tracks/number of total moving tracks x 100; % of bidirectional tracks: number of bidirectional tracks/number of total moving tracks x 100% (Qu et al., 2019; Stepanova et al., 2010).

#### Live imaging of Ca^2+^ influx by GCaMP3S in dopaminergic neurons

Ventral midbrain neurons from GCaMP3S-IRES-DAT-Cre neurons expressing GCaMP3S only in dopaminergic neurons were imaged at 14 DIV. To induce neuronal activity, neurons were washed 1x with complete HBSS, then incubated with HBSS with DMSO control, 20 µM bicuculline to induce neuronal activity, or 20 µM bicuculline + 50 µM D-AP5 + 20 µM CNQX to completely suppress glutamatergic stimulation. To test whether GatastatinG2 altered Ca^2+^ influx, we treated GCaMP3S-IRES-DAT-Cre neurons with DMSO, 20 µM bicuculline, or 20 µM bicuculline + 10 µM GatastatinG2. We recorded movies with a 60× objective at 2 s/frame for 3 min on a Nikon Ti2 Eclipse microscope equipped with a dual-camera Hamamatsu ORCA-Fusion BT C15400, a Yokogawa CSU-W1 spinning disk module, and a Tokai-Hit live imaging incubator, 1 min after treatment. Peak amplitude was measured as the fold change of the ΔF/F_0_ average peak amplitude of control neurons. Fluorescence intensity was determined on a 56 µm^2^ ROI within the cell body.

#### Live imaging of SV release with MT dynamics in dopaminergic neurons

To test the relationship between MT dynamics and synaptic exocytosis via the SYP1-pHluorin assay using electrical field stimulation, we infected DAT-IRES-Cre neurons with flex SYP1-pHluorin, flex SYP1-HaloTag (pan-synaptic marker), and flex EB3-TdTomato at 7-10DIV. Imaging of live neurons was performed at 14-17 DIV in HBSS with 10µM CNQX, 50 µM D-AP5, 5µM Nimodipine, and 10 µM ZD7288 at 37 °C to suppress both evoked and pacemaker spontaneous activity and SV release. Movies were acquired using an IX83 Andor Revolution XD Spinning Disk Confocal System. Confocal live imaging was performed by acquiring 3 stacks of 0.5 µm each, for all three channels at a rate of 1 frame/5 sec. Action potentials (AP) were evoked by passing 1 ms bipolar current at 100 mA through platinum-iridium electrodes connected to a live imaging dish containing imaging buffer at 10 Hz using a digital stimulator (Master8) and a stimulator isolator. HBSS buffer supplemented with 100 mM NH_4_Cl was perfused at the end of the recording to assess maximum pHluorin signal. Peak amplitude was measured as the fold change of ΔF/ΔF_NH4Cl_. For each movie, we divided the fraction of all comet starts that occurred during stimulation by the fraction of the movie spent under stimulation. For the bouton-specific ratio, we divided the fraction of all starts that occurred at boutons during stimulation by the product of the stimulation time fraction and the fraction of neurite area occupied by boutons (bouton count × mean bouton area [1 µm²] / estimated neurite area).

#### Acute brain slice preparation and fast scan cyclic voltammetry (FSCV)

3–4 months old C57BL/6 and α-synuclein KO mice were euthanized by cervical dislocation, and 250 μm-thick 10° angle oblique horizontal striatal slices were prepared on a vibratome (VT1200; Leica, Sloms, Germany) in oxygenated ice-cold ‘cutting’ artificial cerebrospinal fluid (ACSF) containing (in mM): 194 sucrose, 30 NaCl, 4.5 KCl, 26 NaHCO3, 6 MgCl2·6H2O, 1.2 NaH2PO4, and 10 D-glucose (pH 7.4, 290±5 mOsm). Slices were then transferred to oxygenated ‘normal’ ACSF containing (in mM): 125.2 NaCl, 2.5 KCl, 26 NaHCO3, 1.3 MgCl2·6H2O, 2.4 CaCl2, 0.3 NaH2PO4, 0.3 KH2PO4, and 10 D-glucose (pH 7.4, 290±5 mOsm) for 20 min at 34°C and then for 20 min at room temperature for slice recovery. After recovery, brain slices were split into hemispheres and incubated in separate chambers containing either ACSF (control) or GatastatinG2 (10 μM). After 1 hour of incubation, slices were transferred to a recording chamber maintained at 34°C and continuously perfused with their respective incubation solutions, during which dopamine release was measured. The carbon fiber electrode (5 μm diameter) was inserted into the dorsolateral striatum, 25-50 μm below the surface, and a stimulation electrode (WPI, Sarasota, FL) was placed on the surface 50-100 μm away from the carbon fiber electrode. Evoked dopamine release was recorded from 3-5 distinct spots per slice in each condition, with recording times (1, 2, 3, and 4 hours) kept consistent across groups. FSCV recordings were conducted using Igor Pro 6 (Wavemetrics, OR) with a custom-made FSCV macro (written by Dr. Eugene Mosharov, Columbia University/NYPSI). A background triangular-wave voltage (−450 to +800 mV at 294 V/s vs Ag/AgCl) was applied to the electrode every 100 ms. Dopamine peak currents were recorded with Axopatch 200B (Molecular Devices) and digitized with InstruTECH ITC-18 at 10 kHz. To convert the dopamine peak current into dopamine concentration, carbon fiber electrodes were calibrated with 1 µM dopamine (Sigma-Aldrich) before every recording.

#### Crude synaptosomal fraction isolation from striatum

Adult WT mice (3 months old) were euthanized, brains were rapidly extracted, and the striatum was dissected and immediately frozen on dry ice. Samples were weighed and homogenized using a Potter–Elvehjem homogenizer at a 1:10 ratio (mg tissue: µl buffer) in ice-cold synaptosome isolation buffer (5 mM HEPES, 0.32 M sucrose in PBS) supplemented with protease and phosphatase inhibitors (Thermo Fisher, Cat. 1861281). Homogenization was performed with 15 strokes using a Teflon pestle. Homogenates were centrifuged at 1,000 × g for 10 min at 4°C to pellet nuclei and debris (P1). The resulting supernatant (S1) was centrifuged at 12,000 × g for 10 min at 4°C to obtain a crude synaptosomal pellet (P2), containing synaptic membranes and mitochondria, and a supernatant (S2) containing light membranes and soluble protein (Wirths, 2017). Final synaptosomal pellets were lysed in RIPA buffer supplemented with protease and phosphatase inhibitors for 10 min at 4°C, diluted with 5× Laemmli buffer (up to 1x), and boiled for 5 min at 96°C prior to SDS–PAGE.

#### Western blot analyses

Cells were lysed in Laemmli sample buffer and boiled at 96°C for 5 min. Cell lysates were sonicated by a probe sonicator to shear cellular debris and genomic DNA. Proteins were separated by a 4-12% Bis-Tris gel (Invitrogen) and transferred onto a nitrocellulose membrane. After blocking in 5% milk/TBS or BSA/TBS, membranes were incubated with primary antibodies at 4°C overnight, followed by a 1 h incubation with secondary antibodies. Image acquisition was performed with an Odyssey imaging system (LI-COR Biosciences, NE) and analyzed with Odyssey software.

#### Peptide array overlay assay

To map the epitopes within α-synuclein that bind to soluble cromathography-purified porcine α/β-tubulin dimers (Cytoskeleton, Cat# T240-B), human recombinant γ-tubulin (Creative BioMart), or Taxol-stabilized MTs, human α-synuclein protein was synthesized as fragments of 33 linear peptides (with N-terminal acetylation) covalently C-terminally bound to a cellulose membrane. Membranes were generated using the SPOT synthesis technique by JPT Peptide Technologies (Berlin, Germany). Each spot on the membrane contained a 15-amino acid peptide fragment that tiled the entire human α-synuclein sequence, including four peptides to map each mutation, overlapping with the next peptide fragment by 11 amino acids. Spots were arrayed in a grid, 20 per row. For the overlay assay, the membrane was saturated in 3% BSA in TBS, incubated with either soluble tubulin dimers, Taxol-stabilized MTs, or γ-tubulin in 1% BSA in BRB80 (80 mM PIPES, pH 7.2, 1 mM EGTA, 1 mM MgCl2) for 2 h at a final concentration of 2.5 μg/ml at RT. Taxol at a final concentration of 10 µM was added during the incubation of Taxol-stabilized MTs with the peptide array membrane. The membrane was then washed with TBS-T three times and subsequently incubated with the anti-*α*-tubulin antibody (Sigma-Aldrich, Cat# T9026), followed by an IR-conjugated secondary anti-mouse antibody to reveal binding via classical Western blot. Relative binding strength (%) was calculated after subtracting the background and normalizing the integrated density of each spot to the most intense spot in the membrane in each experiment.

#### Enzymatic C-terminal tail cleavage of α/β-tubulin dimers and taxol-stabilized MT preparation

To determine whether α-synuclein binding in the peptide array assay depends on the presence of the C-terminal tails of α- and/or β-tubulin, regions that harbor most tubulin post-translational modifications, we enzymatically removed these tails using Subtilisin A. Lyophilized porcine tubulin (Cytoskeleton, Cat# T240-B) was resuspended on ice in BRB80 supplemented with 1 mM GTP to a final concentration of 6 mg/ml. The tubulin was then incubated with increasing concentrations of Subtilisin A (Sigma-Aldrich, Cat# P5380) at 25°C for 40 min. To assess the extent of C-terminal cleavage, aliquots of the reaction were lysed in Laemmli buffer, resolved on a 10% Bis-Tris gel (Invitrogen), and transferred onto a nitrocellulose membrane. The membrane was probed with an antibody against tyrosinated α-tubulin (YL1/2; Sigma-Aldrich, Cat# MAB1864), which recognizes an epitope located within the last few amino acids of the α-tubulin C-terminal tail. This allowed the detection of uncleaved α-tubulin. Band intensities were normalized to total α-tubulin levels (Proteintech, Cat# 11224). A 1:60 (w/w) enzyme-to-tubulin ratio was selected, as it resulted in complete removal of the C-terminal tails, indicated by the disappearance of the tyrosinated α-tubulin signal (**Fig. S5A**). Following digestion, both subtilisin-treated and control (no enzyme) samples were desalted and cleared of C-terminal fragments and residual enzyme using Zeba MWCO 40 kDa desalting columns (Thermo Fisher, Cat# A57759), collecting the flow-through. All steps were performed at 4°C. Tubulin samples were then subjected to ultracentrifugation at 100,000 × g for 10 minutes at 4°C (Beckman Optima MAX-TL with TLA 100.3 rotor) to remove aggregates. The resulting supernatant, containing soluble tubulin dimers, was either used directly for the peptide array assay or polymerized into Taxol-stabilized MTs. For MT polymerization, full-length or C-terminally cleaved tubulin was diluted to 3 mg/mL in BRB80 supplemented with 1 mM GTP and incubated at 37°C for 10 min. Taxol was then added stepwise (1 to 10 µM), and MTs were pelleted by ultracentrifugation at 100,000 × g at 25°C (Beckman Optima MAX-TL with TLA 100.3 rotor) prior to resuspension in BRB80 containing 1 mM GTP and 10 µM Taxol.

#### Immunofluorescence analysis and AiryScan microscopy

Neurons were fixed in 4% PFA + 4% sucrose for 20 min at RT. Cells were then washed in PBS, permeabilized with 0.2% Triton X-100 for 10 min at RT, blocked in 3% BSA in PBS for 1 h at RT, and stained with primary antibodies for 2 h followed by secondary antibodies for 1 h at RT. Samples were kept in live-cell dishes, mounted using Ibidi mounting medium, and imaged using an IX83 Andor Revolution XD Spinning Disk Confocal System. The microscope was equipped with a 60x 3 /1.30 silicon UPlanSApo objective and a multi-axis stage controller (ASI MS-2000). Movies were acquired with an Andor iXon Ultra EMCCD camera and Andor iQ 3.6.2 imaging software. All images were analyzed using ImageJ. Analysis of colocalization (**Fig. 2B**) was performed by measuring colocalization between α-synuclein and γ-tubulin after creating a single mask on tyrosine hydroxylase (TH) signal prior to using the JaCOP colocalization plugin. Spatial cross-correlation (**Fig. S2A**) was assessed by translating the γ-tubulin channel by 15 pixels along the X-axis relative to synapsin, revealing a significant reduction in Manders’ coefficients toward a chance-level plateau. This indicates that the observed colocalization is not attributable to random spatial overlap. To measure the number of dopaminergic synapses containing either α-synuclein or γ-tubulin signal, or both (**Fig. 4G, H**), confocal images of endogenous α-synuclein and γ-tubulin staining were overlaid onto a mask created from the TH signal, used to isolate dopaminergic neurons. Synaptic puncta were segmented based on synapsin1 signal and filtered by area (0.2–2 µm²). γ-Tubulin and α-synuclein positivity were defined as a contiguous signal area ≥16 pixels within each synaptic ROI. Thresholds were calibrated once based on a visually detectable signal and kept constant across all images. Quantification was performed across 9 fields of view from 3 independent images, for a total of 307 synapses. Qualitative Airyscan images of the same immunostainings (**Fig. 4F**) were acquired using a Zeiss LSM 800 confocal microscope equipped with an Airyscan module, using a 63x objective (Plan-Apochromat, NA 1.4). Single-stack (0.2µm thick) images were obtained and processed for super-resolution using Zen Blue 2.1 software. Manders’ coefficients between synapsin1 and α-synuclein, or between synapsin1 and γ-tubulin, were computed on the same whole-field confocal images, restricted to TH-gated regions, providing a threshold-independent measure of synaptic association (**Fig. S4F)**. This global analysis complements the per-synapse classification reported in the main (**Fig. 4F, G)**. For post-imaging analyses (**Fig. 6G, H**), movies were fixed immediately after live imaging and processed for immunostaining under identical conditions across all samples. Quantification of pS129-α-syn puncta was performed on the same synapses previously analyzed during live imaging.

#### Proximity Ligation Assay (PLA)

After fixation, permeabilization, and saturation as described for IF above, samples were incubated with primary antibodies against α-synuclein and γ-tubulin or pS129 α-synuclein and γ-tubulin for 2 h at 37°C, followed by 1 h ligase incubation and 2 h incubation with polymerase as described in the DuoLink protocol by the vendor (Sigma-Aldrich). Samples were acquired using an IX83 Andor Revolution XD Spinning Disk Confocal System equipped as mentioned above. Analysis of synaptic PLA puncta density (**Fig. S6D**) was performed by counting the number of PLA puncta overlapping with the synapsin signal and dividing it by the area covered by synapses in that image.

#### Molecular docking and molecular dynamics simulations

Molecular docking between the α/β-tubulin dimer and peptide 33 of α-synuclein was performed using AlphaFold3 with a fully blind prediction strategy, generating five candidate complexes (Abramson et al., 2024). GDP and GTP ligands were included in the simulations according to the experimental tubulin structure (PDB: 1TUB). Predicted binding affinities were evaluated using PRODIGY (Xue et al., 2016). All docking poses were subjected to 500 ns molecular dynamics (MD) simulations under periodic boundary conditions using the CHARMM36 force field, with ligand parameters generated by SwissParam. After steepest-descent minimization and equilibration, simulations were run under NPT conditions at 300 K and 1 bar, using the V-rescale thermostat and Parrinello–Rahman barostat. Hydrogen-containing bonds were constrained with LINCS, enabling a 2 fs timestep, and electrostatics were treated using Particle Mesh Ewald (PME). Binding stability during MD simulations was evaluated by measuring the distance between the geometric centers of the peptide and the binding site every 1 ns, with the binding site defined as residues within 12 Å of the peptide in the initial docking configuration.

## QUANTIFICATION AND STATISTICAL ANALYSIS

Data are shown as means ± SEMs from at least 3 independent experiments, and graphs were generated with GraphPad Prism software. Immunofluorescence, time-lapse movie, and western blot image analysis were performed with ImageJ (Fiji). Statistical analysis between two groups was performed using the Mann-Whitney non-parametric test (**Fig. 1H, 4B; Fig. S2A, S2C, S3C, S4A, S4B, S6D**). Comparison among 3 or more groups was performed using the Kruskal-Wallis test with Dunn’s multiple comparisons test for non-parametric unpaired one-way ANOVA (unpaired samples N < 15 in **Fig. 1C, 1G, 1K, 2G, 3B, 3E, 4E, 4F, 5J, 6E, 6G, 7F, 7G; Fig. S1A, S1B, S2D, S3A, S4C, S4D, S5B, S6B, S6C, S7B**). For FSCV, all data were analyzed using Igor Pro. We used two-way ANOVA to compare each time point (**Fig. 3I**) and a paired t-test in **Fig. 3J** and **S3B**. Statistical significance was set at p < 0.05. No statistical analysis was performed on the pep-spot array quantification because binding to tubulin dimers and γ-tubulin was repeated twice (n=2), whereas binding to MTs was assessed in a single experiment (n=1).

## ACKNOWLEGMENTS

This project was funded by a TIGER Grant from the Taub Institute for Research on Alzheimer’s Disease and the Aging Brain at Columbia University to F.B., a Stabilization Fund from Columbia University to F.B., an R01AG050658 (NIH/NIA) award to F.B. and a Postdoctoral Fellowship Award from the Parkinson’s Foundation PF-PRF-1585486 to A.C. This study was also funded by an R01DA07418 and R01NS147157 (NIH) and support from the Freedom Together Foundation to D.S., a European Union’s Horizon 2020 research and innovation program H2020-MSCAITN-2019_EJD: Marie Curie Innovation Training Networks-Grant Agreement No:860070-TubInTrain to G.C. We are grateful to Gregg G. Gundersen for stimulating discussions and access to his microscopes, and to Ryan Dosumu-Johnson for help with viral approaches and live-cell imaging techniques.

## AUTHOR CONTRIBUTIONS

F.B. and A.C. conceptualized the study, designed the experiments, and crafted the manuscript. A.C. performed most of the experiments and analyzed the data. E.K. routinely generated ventral midbrain cultures. S.C. and E.V.M. conducted and analyzed the FSCV experiments while E.M. conducted molecular modeling simulations and analyses. F.B., D.S., G.C., and G.R. provided financial support and supervised data interpretation and experimental strategies. All authors reviewed and approved the final manuscript.

## KEY RESOURCES TABLE

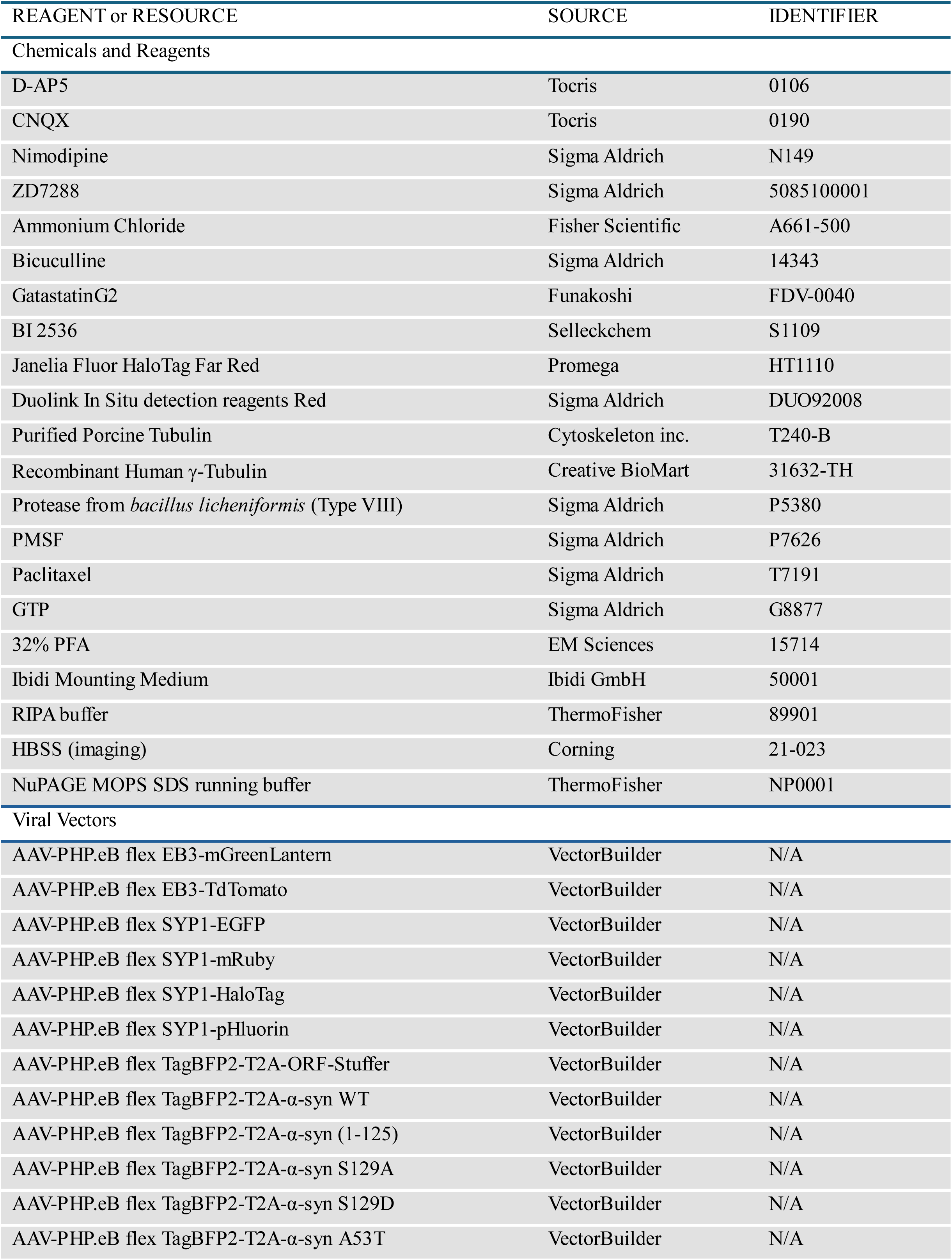
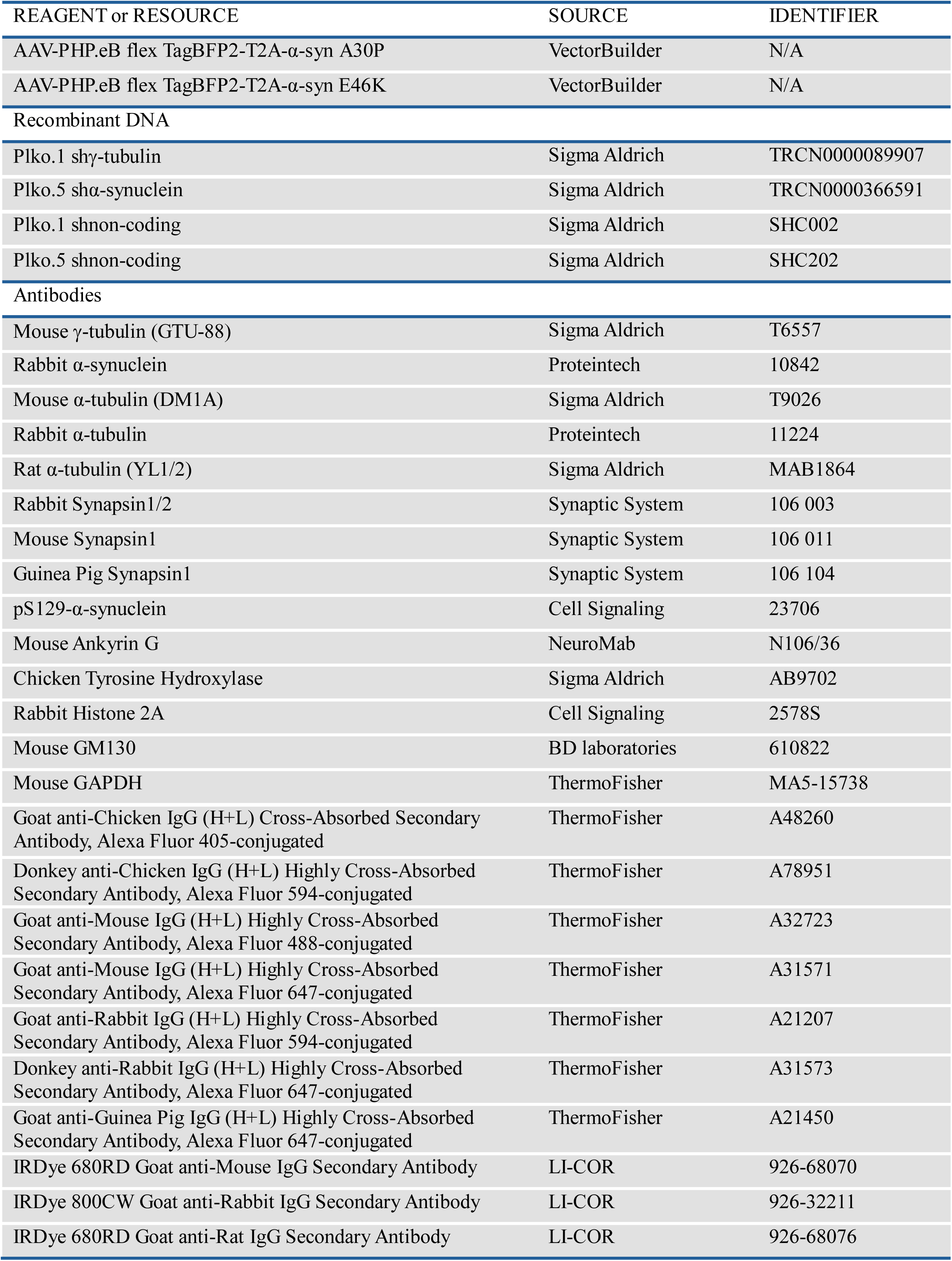
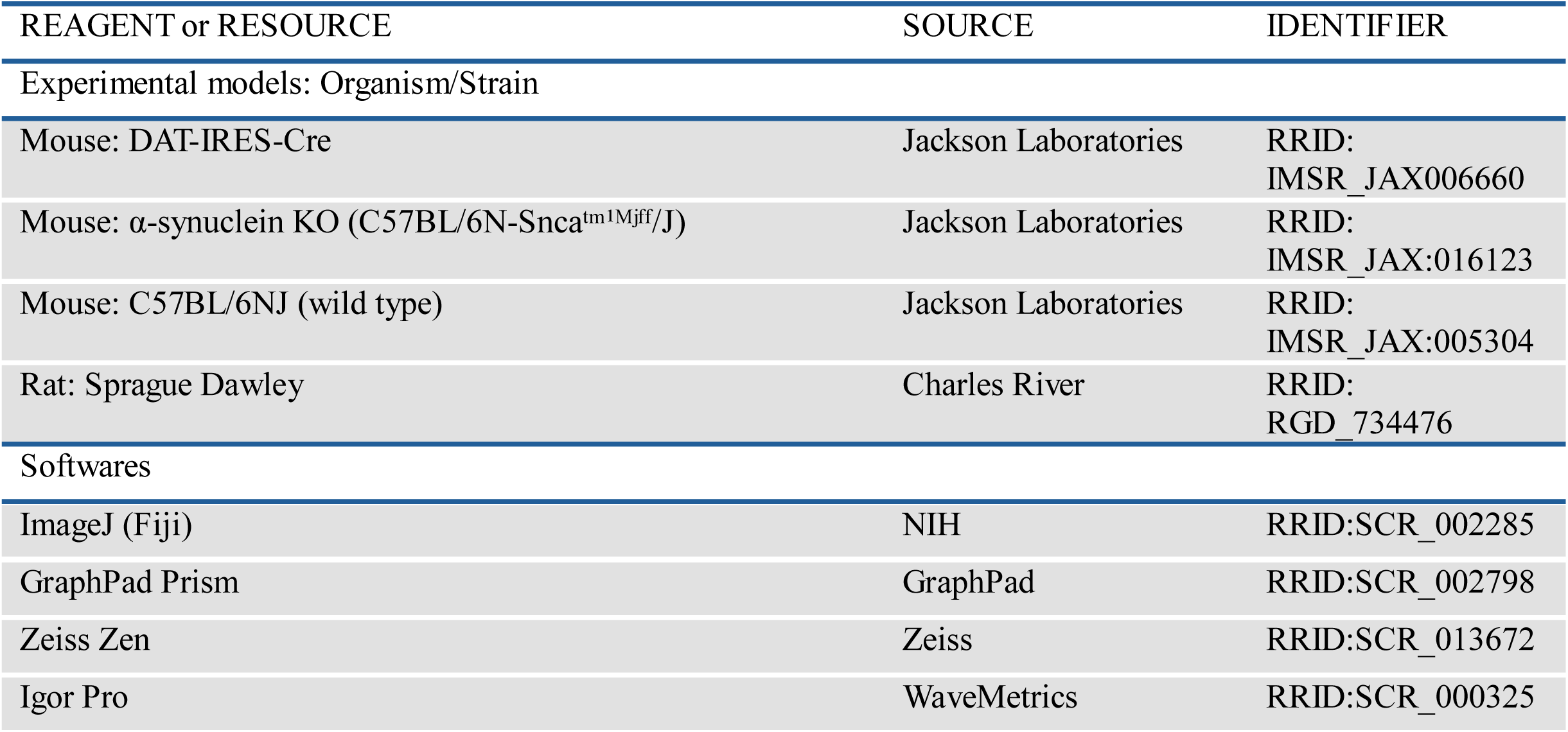

| REAGENT or RESOURCE | SOURCE | IDENTIFIER |
|---|---|---|
| Chemicals and Reagents |  |  |
| D-AP5 | Tocris | 0106 |
| CNQX | Tocris | 0190 |
| Nimodipine | Sigma Aldrich | N149 |
| ZD7288 | Sigma Aldrich | 5085100001 |
| Ammonium Chloride | Fisher Scientific | A661-500 |
| Bicuculline | Sigma Aldrich | 14343 |
| GatastatinG2 | Funakoshi | FDV-0040 |
| BI 2536 | Selleckchem | S1109 |
| Janelia Fluor HaloTag Far Red | Promega | HT1110 |
| Duolink In Situ detection reagents Red | Sigma Aldrich | DUO92008 |
| Purified Porcine Tubulin | Cytoskeleton inc. | T240-B |
| Recombinant Human γ-Tubulin | Creative BioMart | 31632-TH |
| Protease from *bacillus licheniformis* (Type VIII) | Sigma Aldrich | P5380 |
| PMSF | Sigma Aldrich | P7626 |
| Paclitaxel | Sigma Aldrich | T7191 |
| GTP | Sigma Aldrich | G8877 |
| 32% PFA | EM Sciences | 15714 |
| Ibidi Mounting Medium | Ibidi GmbH | 50001 |
| RIPA buffer | ThermoFisher | 89901 |
| HBSS (imaging) | Corning | 21-023 |
| NuPAGE MOPS SDS running buffer | ThermoFisher | NP0001 |
| Viral Vectors |  |  |
| AAV-PHP.eB flex EB3-mGreenLantern | VectorBuilder | N/A |
| AAV-PHP.eB flex EB3-TdTomato | VectorBuilder | N/A |
| AAV-PHP.eB flex SYP1-EGFP | VectorBuilder | N/A |
| AAV-PHP.eB flex SYP1-mRuby | VectorBuilder | N/A |
| AAV-PHP.eB flex SYP1-HaloTag | VectorBuilder | N/A |
| AAV-PHP.eB flex SYP1-pHluorin | VectorBuilder | N/A |
| AAV-PHP.eB flex TagBFP2-T2A-ORF-Stuffer | VectorBuilder | N/A |
| AAV-PHP.eB flex TagBFP2-T2A-α-syn WT | VectorBuilder | N/A |
| AAV-PHP.eB flex TagBFP2-T2A-α-syn (1-125) | VectorBuilder | N/A |
| AAV-PHP.eB flex TagBFP2-T2A-α-syn S129A | VectorBuilder | N/A |
| AAV-PHP.eB flex TagBFP2-T2A-α-syn S129D | VectorBuilder | N/A |
| AAV-PHP.eB flex TagBFP2-T2A-α-syn A53T | VectorBuilder | N/A |
| REAGENT or RESOURCE | SOURCE | IDENTIFIER |
| AAV-PHP.eB flex TagBFP2-T2A-α-syn A30P | VectorBuilder | N/A |
| AAV-PHP.eB flex TagBFP2-T2A-α-syn E46K | VectorBuilder | N/A |
| Recombinant DNA |  |  |
| Plko.1 shγ-tubulin | Sigma Aldrich | TRCN0000089907 |
| Plko.5 shα-synuclein | Sigma Aldrich | TRCN0000366591 |
| Plko.1 shnon-coding | Sigma Aldrich | SHC002 |
| Plko.5 shnon-coding | Sigma Aldrich | SHC202 |
| Antibodies |  |  |
| Mouse γ-tubulin (GTU-88) | Sigma Aldrich | T6557 |
| Rabbit α-synuclein | Proteintech | 10842 |
| Mouse α-tubulin (DM1A) | Sigma Aldrich | T9026 |
| Rabbit α-tubulin | Proteintech | 11224 |
| Rat α-tubulin (YL1/2) | Sigma Aldrich | MAB1864 |
| Rabbit Synapsin1/2 | Synaptic System | 106 003 |
| Mouse Synapsin1 | Synaptic System | 106 011 |
| Guinea Pig Synapsin1 | Synaptic System | 106 104 |
| pS129-α-synuclein | Cell Signaling | 23706 |
| Mouse Ankyrin G | NeuroMab | N106/36 |
| Chicken Tyrosine Hydroxylase | Sigma Aldrich | AB9702 |
| Rabbit Histone 2A | Cell Signaling | 2578S |
| Mouse GM130 | BD laboratories | 610822 |
| Mouse GAPDH | ThermoFisher | MA5-15738 |
| Goat anti-Chicken IgG (H+L) Cross-Absorbed Secondary Antibody, Alexa Fluor 405-conjugated | ThermoFisher | A48260 |
| Donkey anti-Chicken IgG (H+L) Highly Cross-Absorbed Secondary Antibody, Alexa Fluor 594-conjugated | ThermoFisher | A78951 |
| Goat anti-Mouse IgG (H+L) Highly Cross-Absorbed Secondary Antibody, Alexa Fluor 488-conjugated | ThermoFisher | A32723 |
| Goat anti-Mouse IgG (H+L) Highly Cross-Absorbed Secondary Antibody, Alexa Fluor 647-conjugated | ThermoFisher | A31571 |
| Goat anti-Rabbit IgG (H+L) Highly Cross-Absorbed Secondary Antibody, Alexa Fluor 594-conjugated | ThermoFisher | A21207 |
| Donkey anti-Rabbit IgG (H+L) Highly Cross-Absorbed Secondary Antibody, Alexa Fluor 647-conjugated | ThermoFisher | A31573 |
| Goat anti-Guinea Pig IgG (H+L) Highly Cross-Absorbed Secondary Antibody, Alexa Fluor 647-conjugated | ThermoFisher | A21450 |
| IRDye 680RD Goat anti-Mouse IgG Secondary Antibody | LI-COR | 926-68070 |
| IRDye 800CW Goat anti-Rabbit IgG Secondary Antibody | LI-COR | 926-32211 |
| IRDye 680RD Goat anti-Rat IgG Secondary Antibody | LI-COR | 926-68076 |
| REAGENT or RESOURCE | SOURCE | IDENTIFIER |
| Experimental models: Organism/Strain |  |  |
| Mouse: DAT-IRES-Cre | Jackson Laboratories | RRID: IMSR_JAX006660 |
| Mouse: α-synuclein KO (C57BL/6N-Snca<sup>tm1Mjff</sup>/J) | Jackson Laboratories | RRID: IMSR_JAX:016123 |
| Mouse: C57BL/6NJ (wild type) | Jackson Laboratories | RRID: IMSR_JAX:005304 |
| Rat: Sprague Dawley | Charles River | RRID: RGD_734476 |
| Softwares |  |  |
| ImageJ (Fiji) | NIH | RRID:SCR_002285 |
| GraphPad Prism | GraphPad | RRID:SCR_002798 |
| Zeiss Zen | Zeiss | RRID:SCR_013672 |
| Igor Pro | WaveMetrics | RRID:SCR_000325 |

