## SUPPLEMENTARY MATERIAL for "α-Synuclein and γ-Tubulin Cooperatively Regulate Activity-Evoked Presynaptic Microtubule Nucleation to Gate Dopamine Release"

# S1

**A** Additional microtubule dynamics parameters in axons and dendrites of dopaminergic neurons

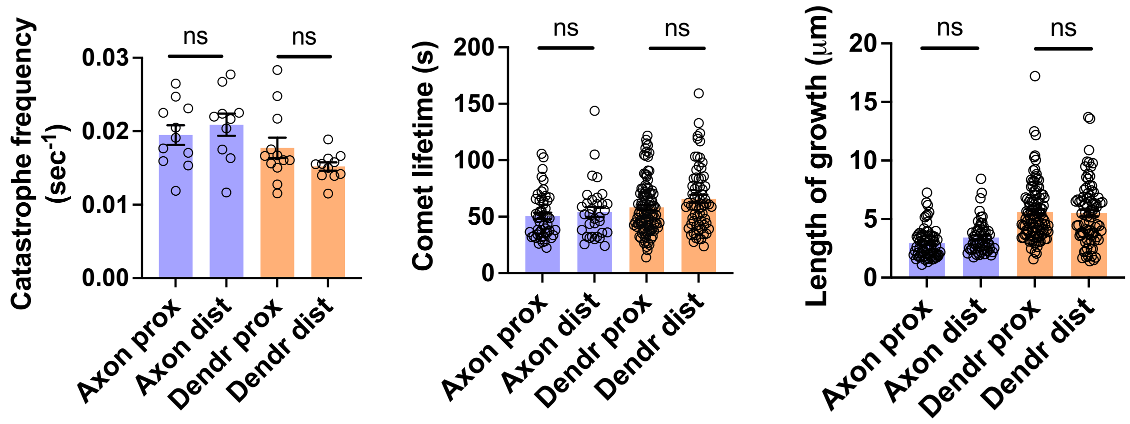


**B** Axonal microtubule dynamics during activity with or without glutamate receptor blockade

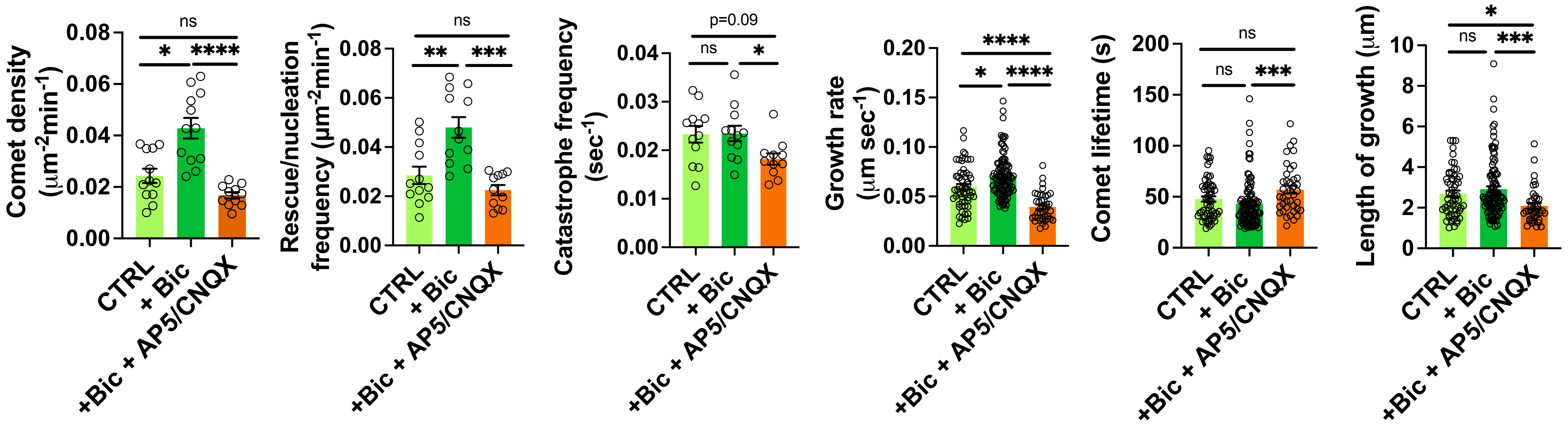

# S2

**A** Cross-correlation analysis of γ-tubulin colocalization with synapsin1/2 in dopaminergic neurons

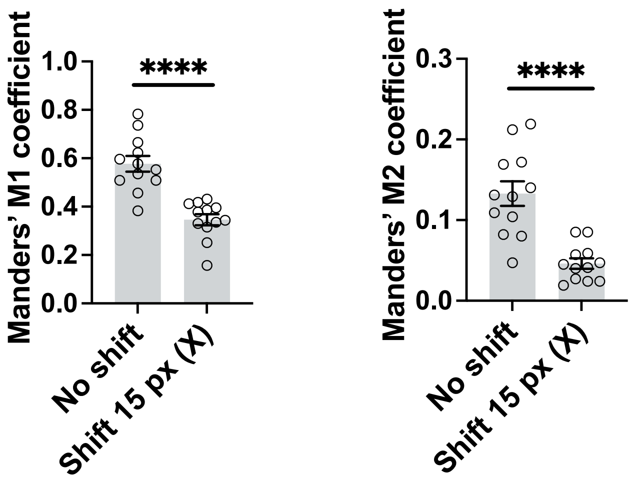


**B** Assessment of crude synaptosomal fractions purity and loading control

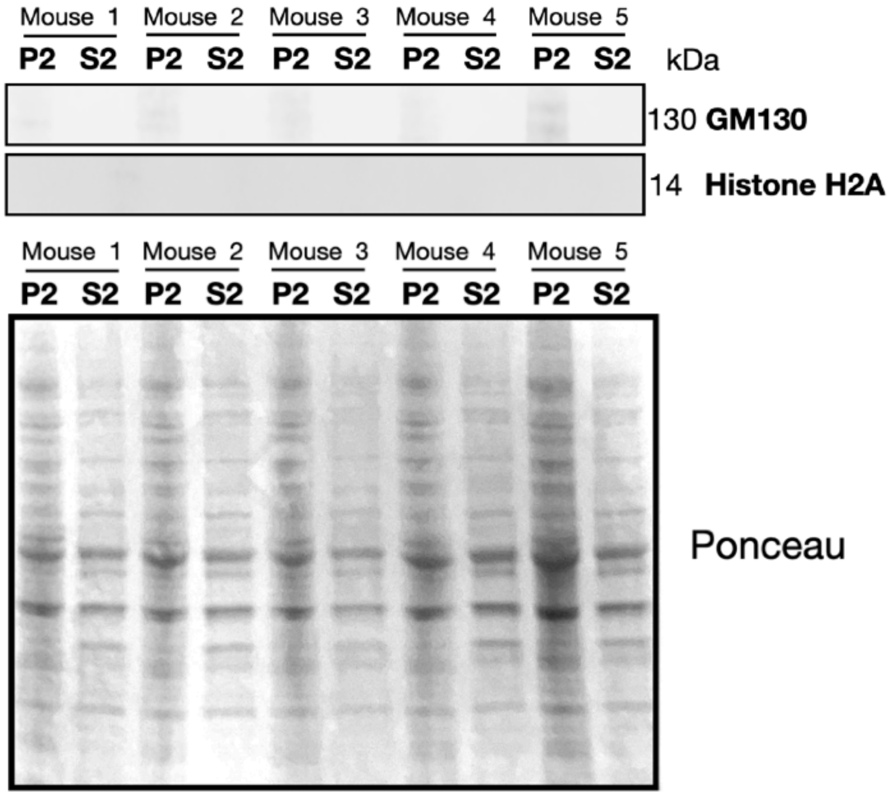


**C** γ-Tubulin levels and distribution at synapses in dopaminergic neurons upon γ-tubulin knockdown

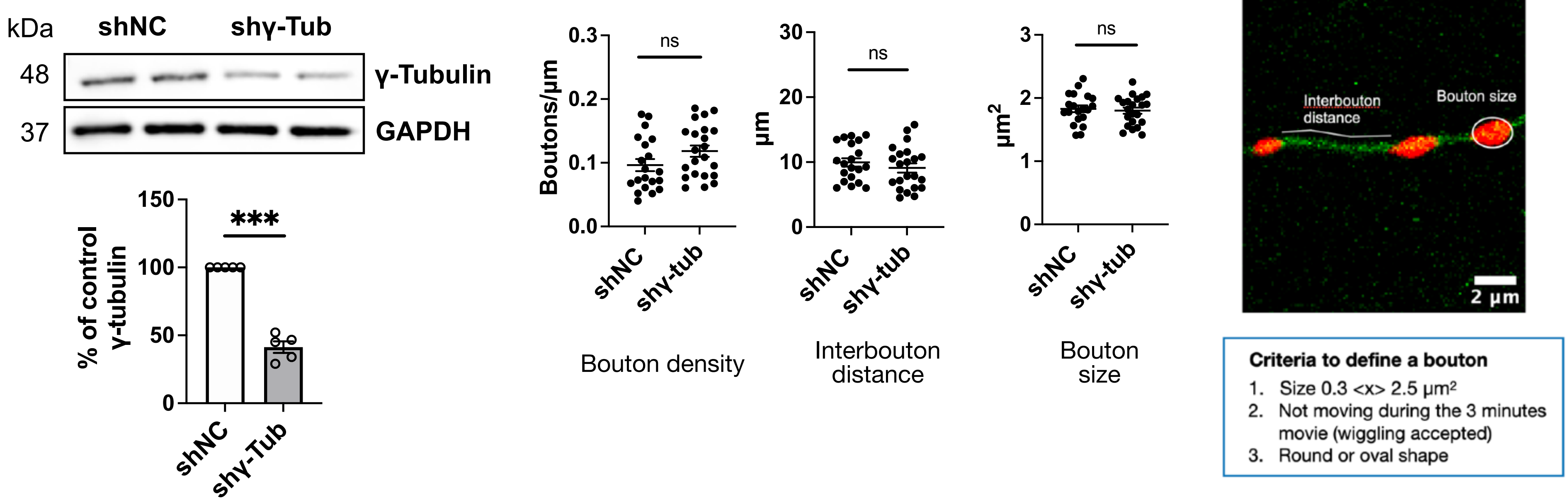


**D** Additional microtubule dynamics parameters and microtubule comet density related to SYP1⁺ puncta in dopaminergic axons upon γ-tubulin knockdown

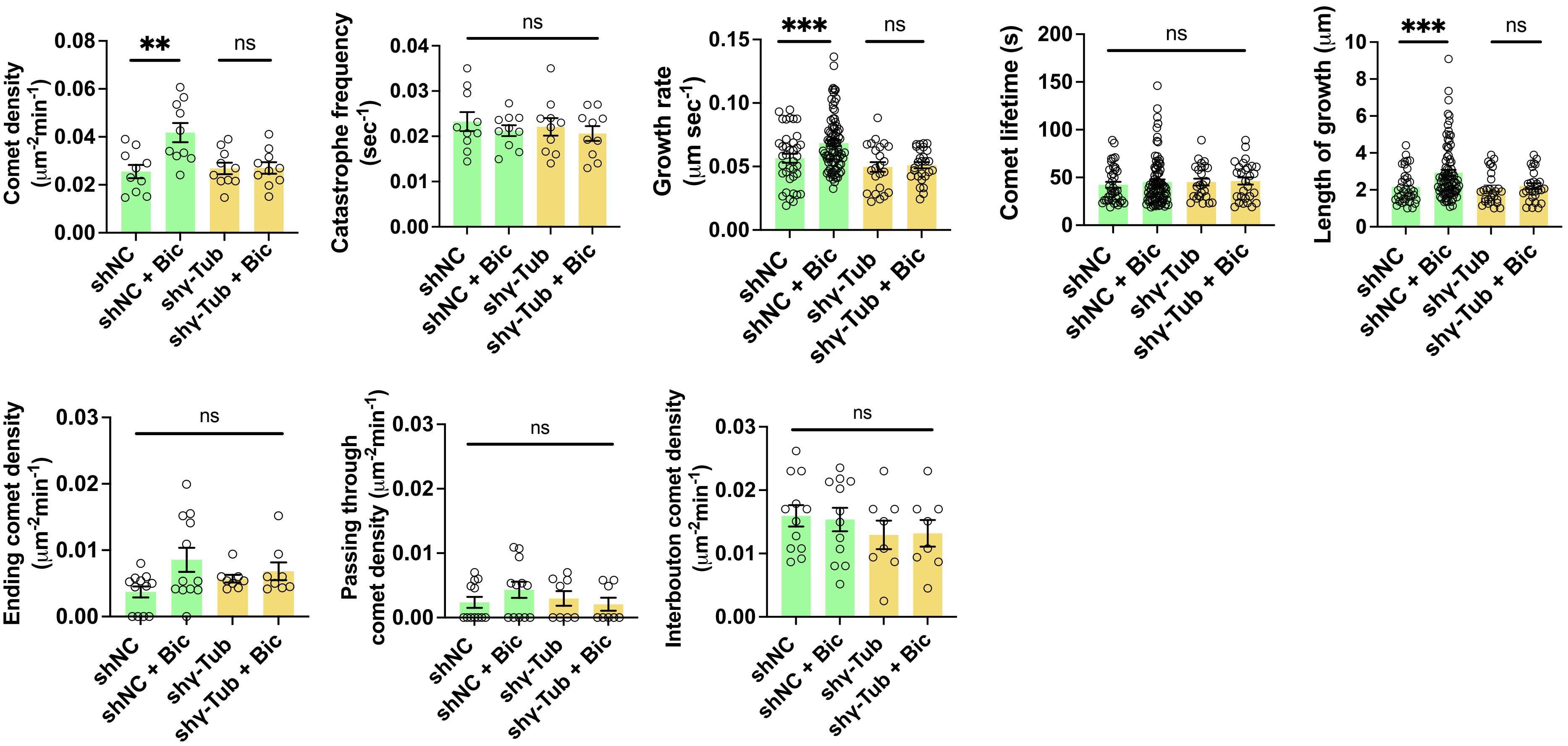

S3

**A** SYP1 puncta motility, starting and ending relative to boutons (in either direction) in dopaminergic axons upon γ-tubulin knockdown or acute gatastatin treatment

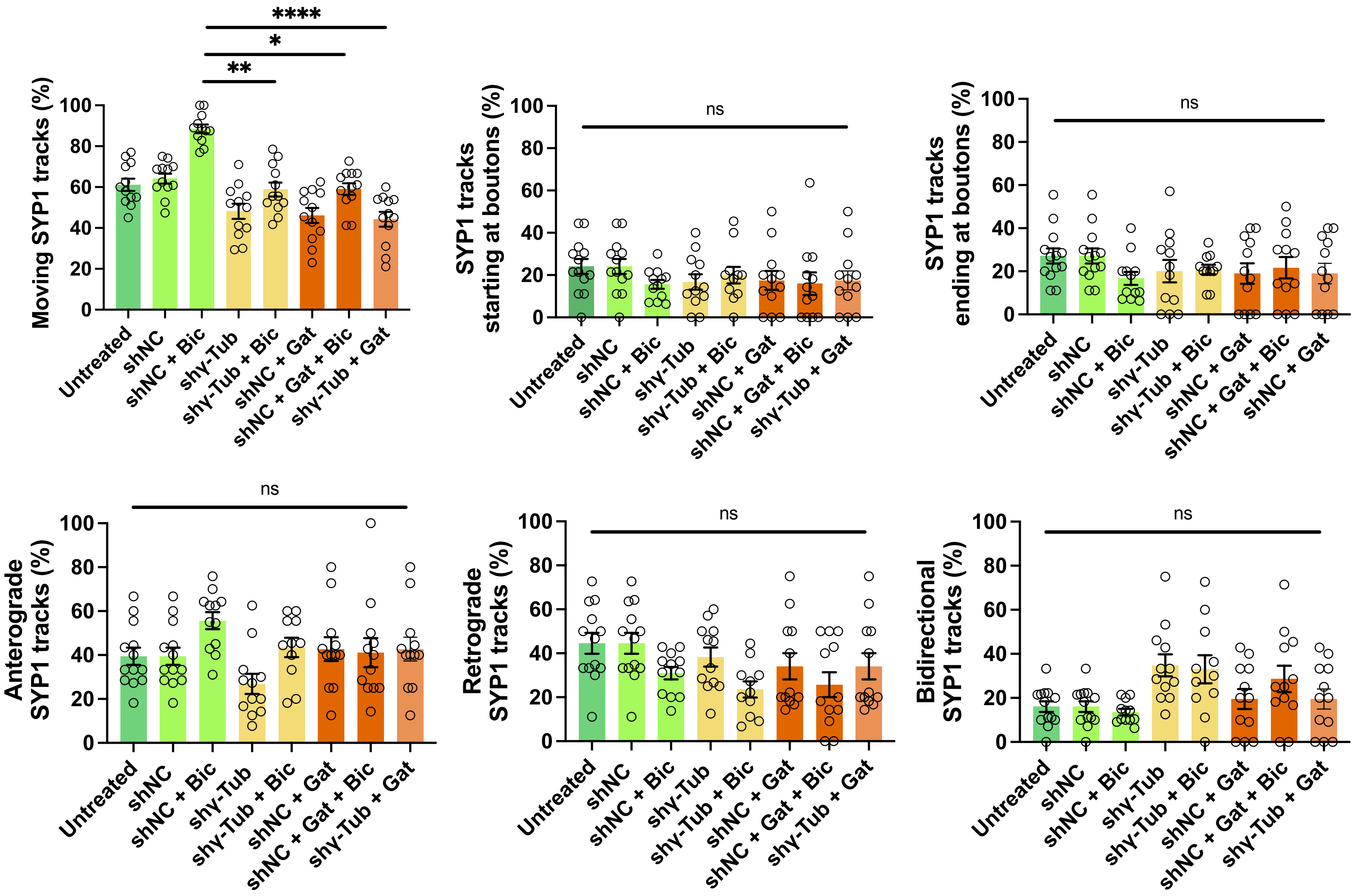


**B** Dopamine half-life ($t_{1/2}$) following 2 to 3 h incubation of GatastatinG2 in WT or α-synuclein KO mice after GatastatinG2 treatment.

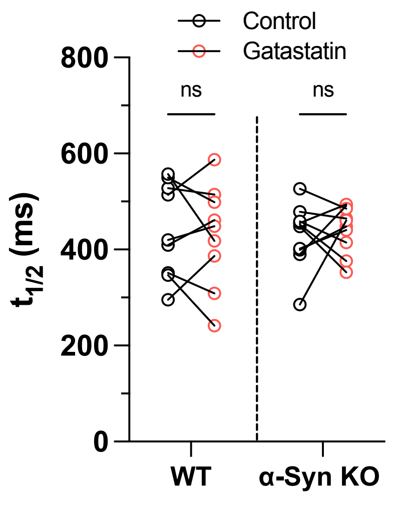


**C** Representative traces and quantification of dopamine release evoked with 5 pulses at 40Hz in WT and α-synuclein KO striatal slices

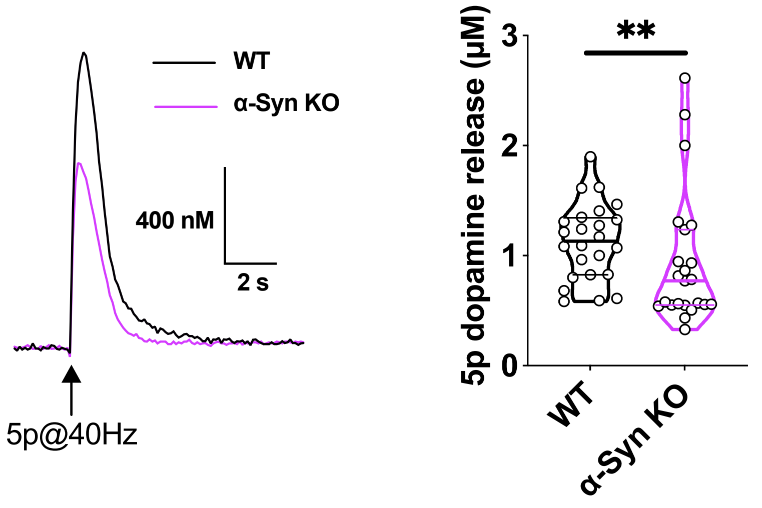

# S4

**A** α-Synuclein levels and distribution at synapses in dopaminergic neurons upon α-synuclein knockdown

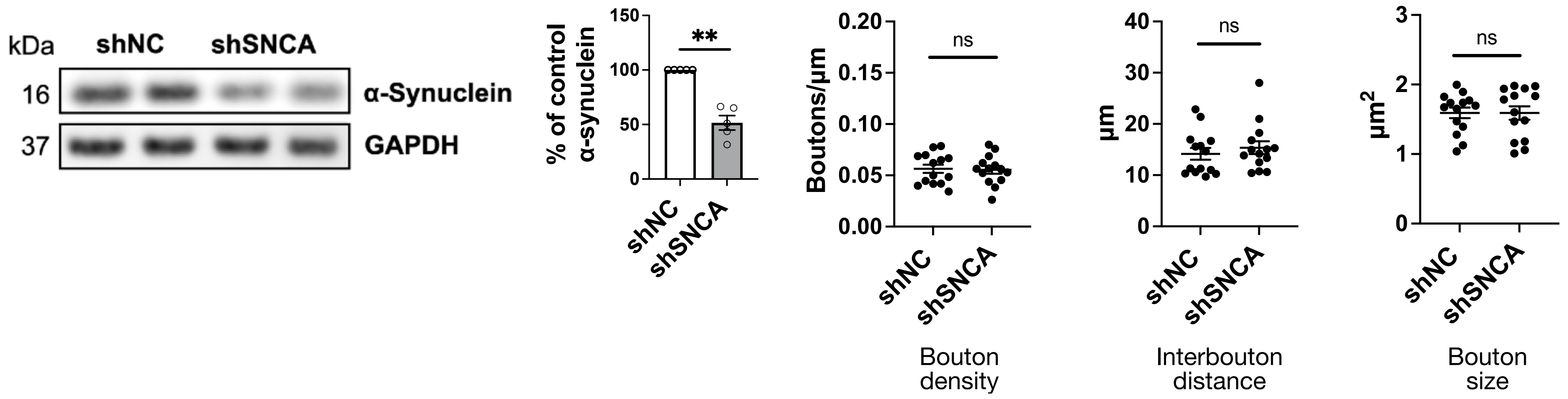


**B** Representative images, Manders' coefficients of colocalization between γ-tubulin and synapsin 1/2, and mean gray value of γ-tubulin upon α-synuclein knockdown

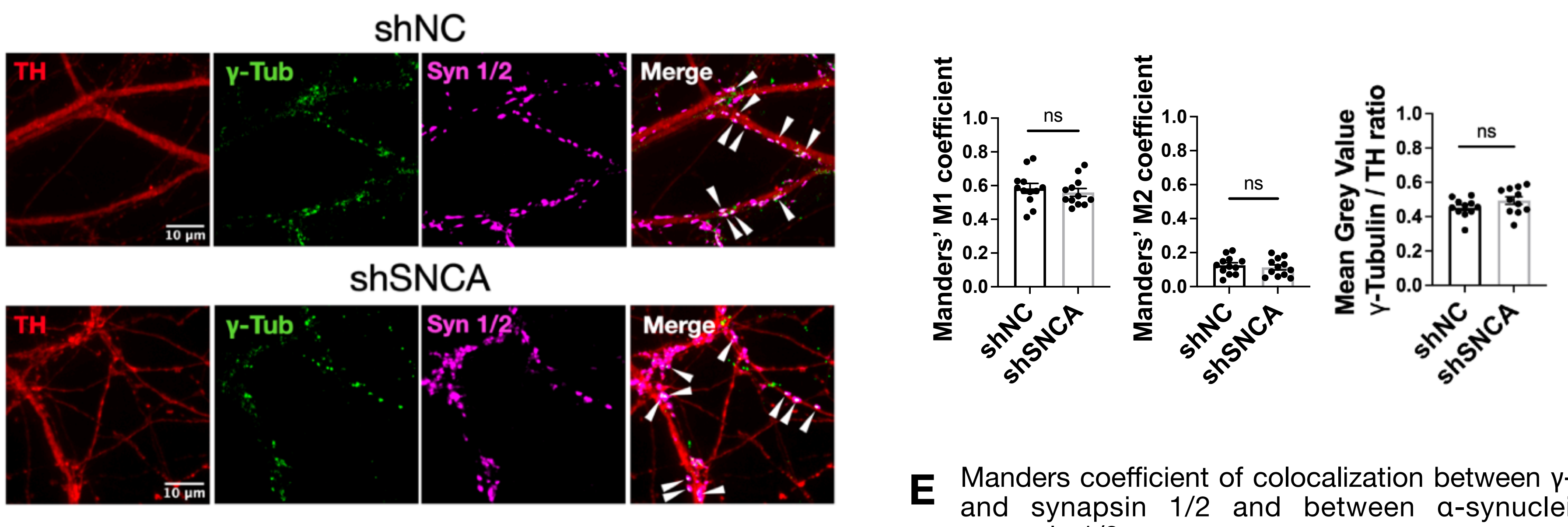


**C** Additional microtubule dynamics parameters and microtubule comet density related to SYP1⁺ puncta in dopaminergic axons upon α-synuclein knockdown

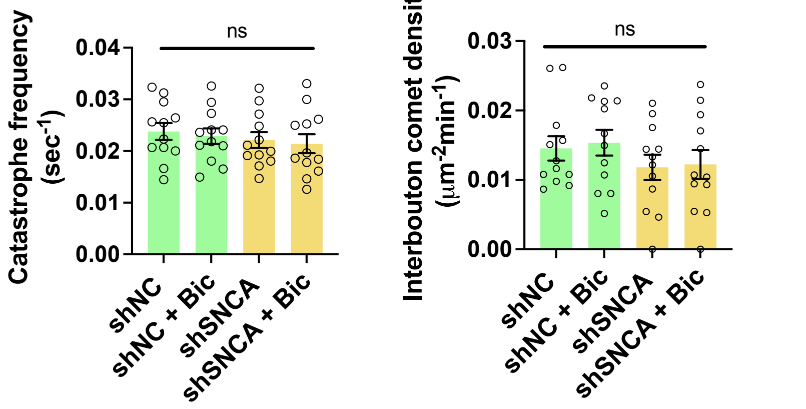


**E** Manders coefficient of colocalization between γ-tubulin and synapsin 1/2 and between α-synuclein and synapsin 1/2

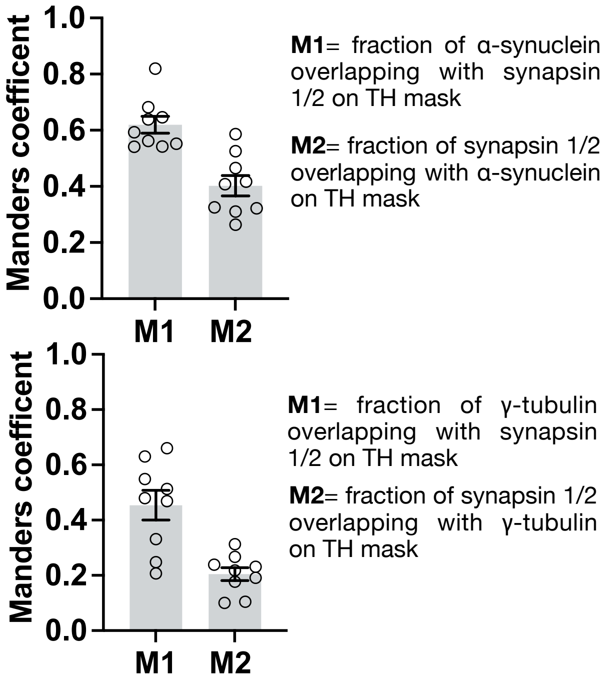


**D** SYP1 puncta motility, starting & ending, only starting or only ending relative to boutons (in either direction) in dopaminergic axons upon α-synuclein knockdown

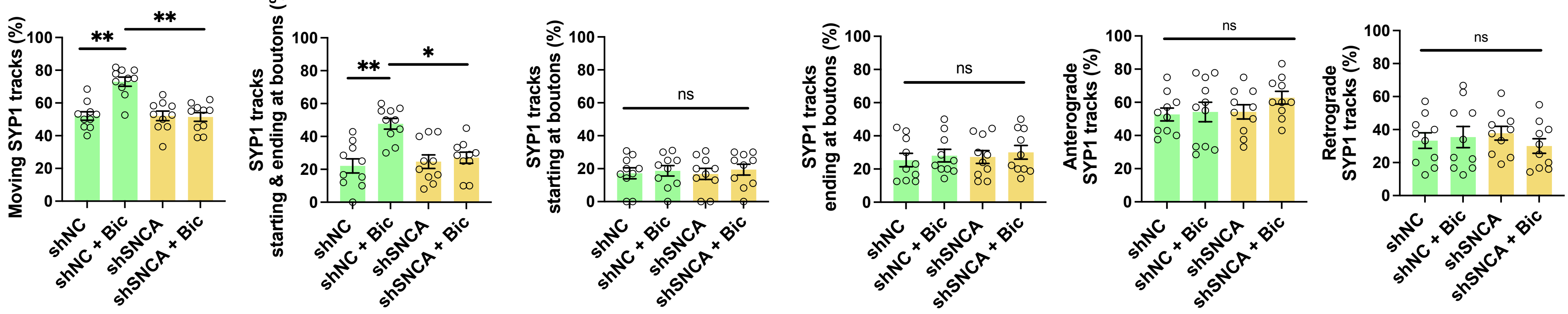

S5

**A** Pep-spot mapping of full-length WT α-synuclein incubated with α/β-tubulin heterodimers after enzymatic removal of α/β-tubulin C-terminal tails or with taxol-stabilized microtubules

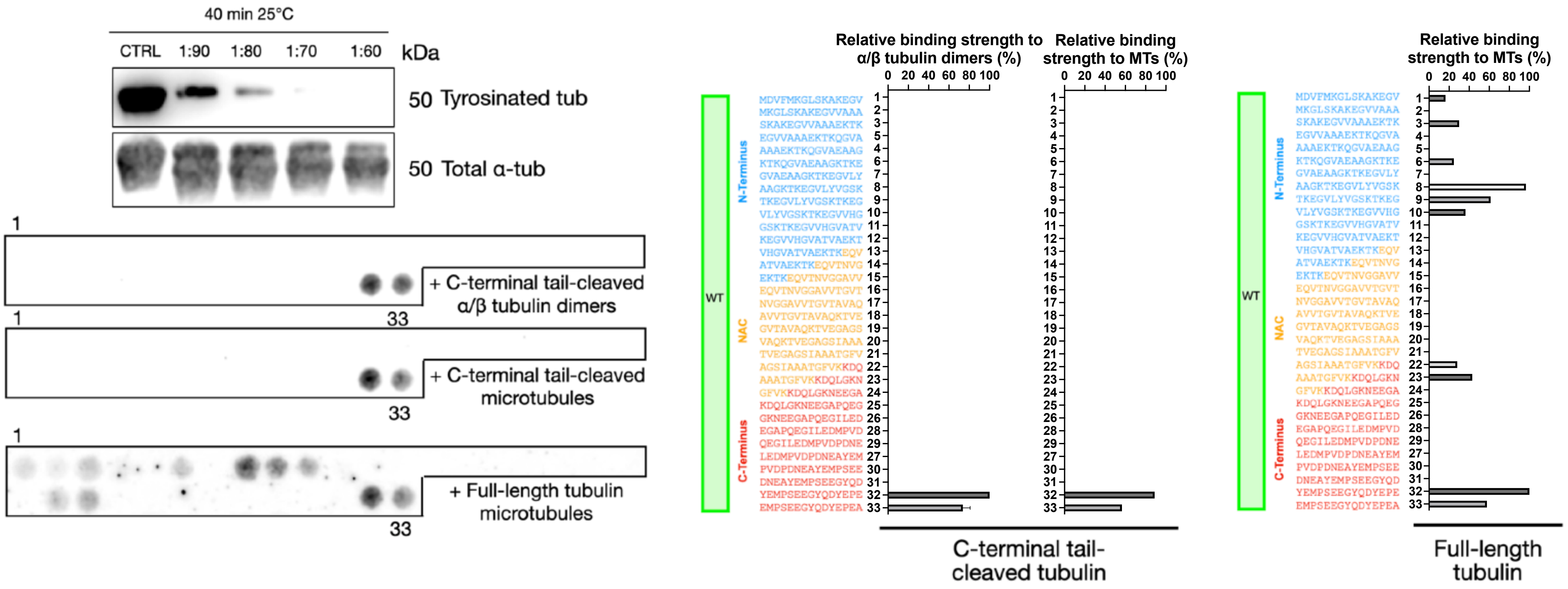


**B** Additional microtubule dynamics parameters and comet density at SYP1$^+$ boutons in dopaminergic axons after α-synuclein knockdown rescued with either ORF-Stuffer control, WT, or C-terminal–truncated (1–125) α-synuclein

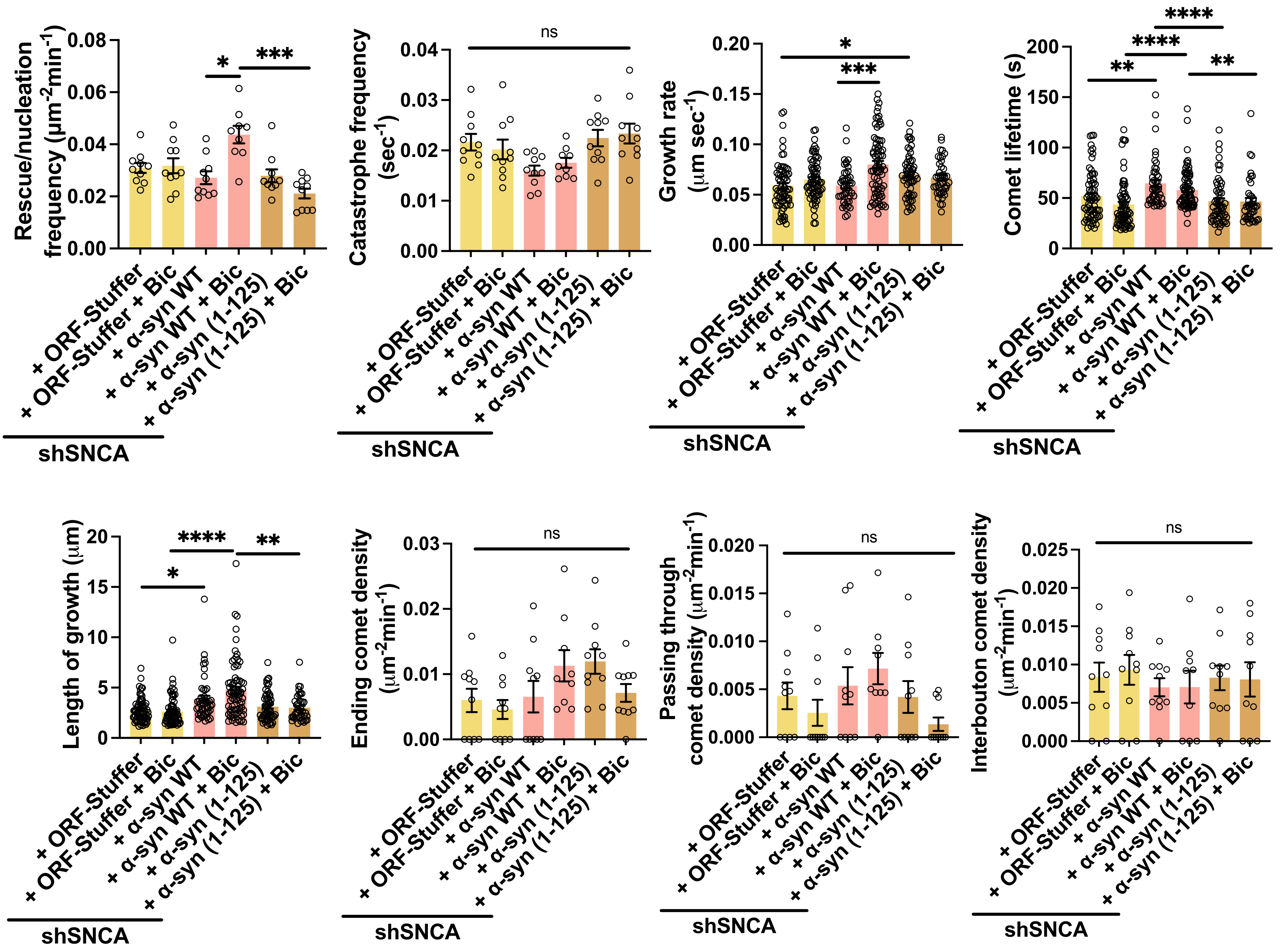

# S6

**A** Pep-spot mapping of WT, S129A, and S129D α-synuclein (AA 126–140, C-terminus) incubated with tubulin dimers after C-terminal tail removal and with taxol-stabilized microtubules

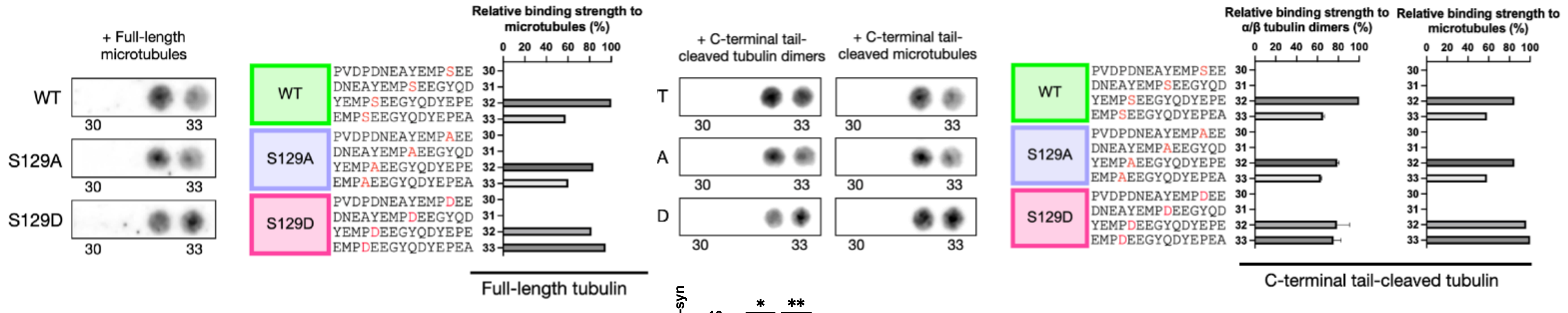


**B** pS129 α-synuclein levels at SYP1⁺ boutons in DAT-Cre neurons after 2h incubation with Bicuculline or 2h BI 2536 PLK2 inhibitor + Bicuculline

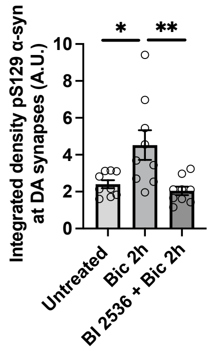


**C** Additional microtubule dynamics parameters and comet density at SYP1⁺ boutons in dopaminergic axons in control (shNC + ORF-Stuffer) or after α-synuclein knockdown and rescue with S129D (phospho-mimetic), S129A (phospho-null) α-synuclein, or control incubated with PLK2 inhibitor BI 2536 (shNC + BI 2536 + ORF-Stuffer)

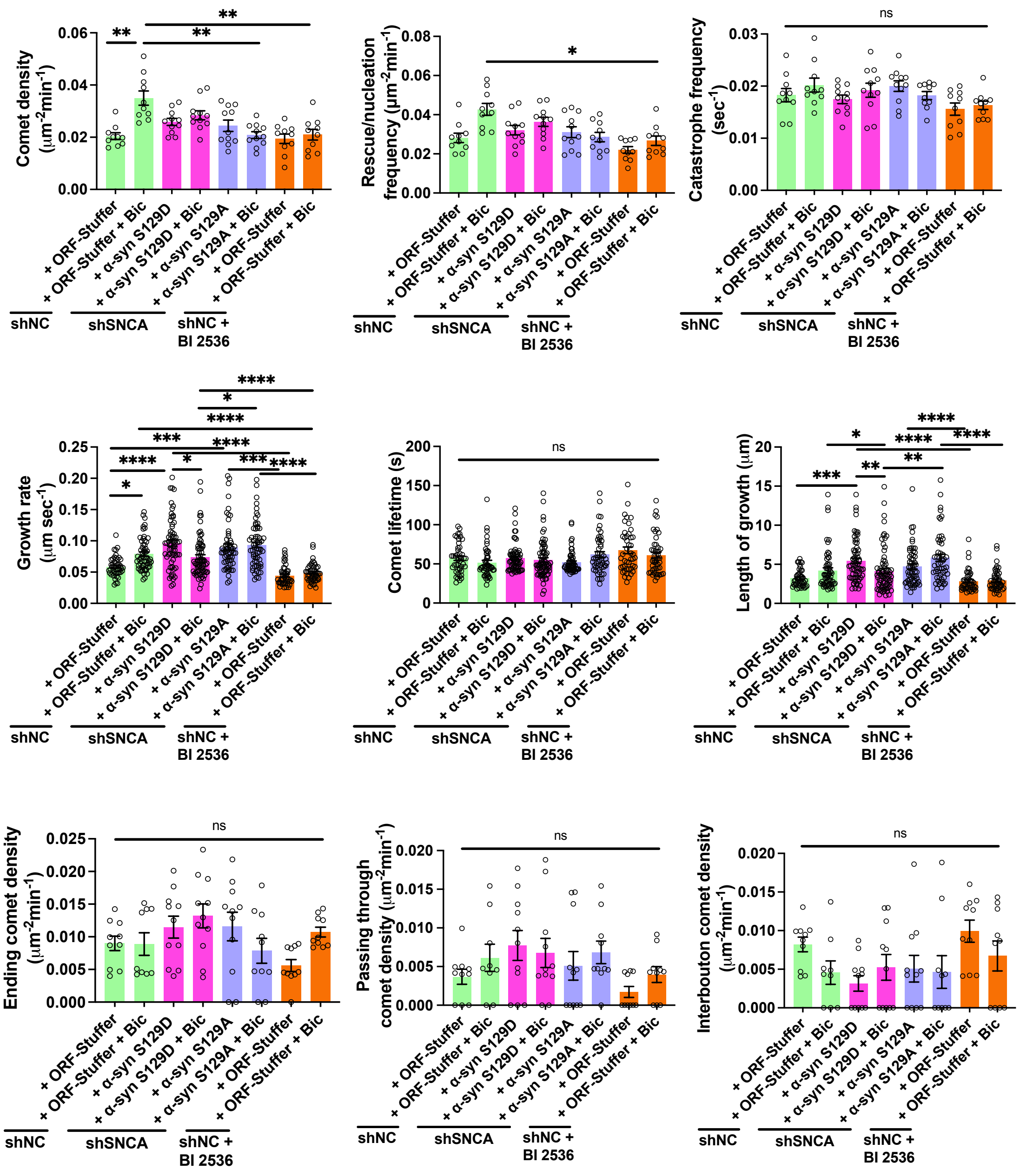


**D** Representative images and synaptic PLA puncta density of PLA between either α-synuclein or pS129-α-synuclein and γ-tubulin with or without neuronal activation

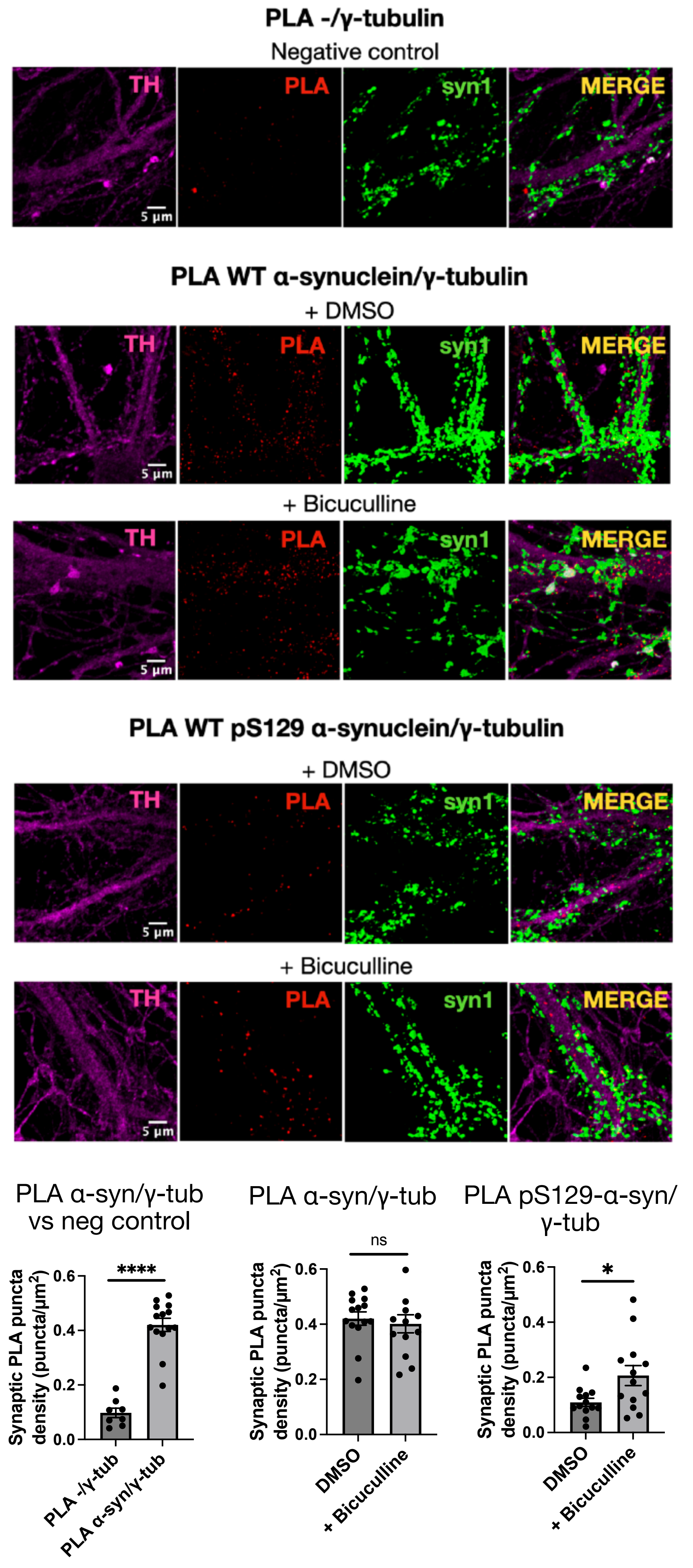

# S7

**A** Pep-spot mapping of WT, A53T, A30P, and E46K α-synuclein incubated with taxol-stabilized microtubules, tubulin dimers after C-terminal tail removal, or taxol-stabilized microtubules generated from C-terminal tail-cleaved tubulin

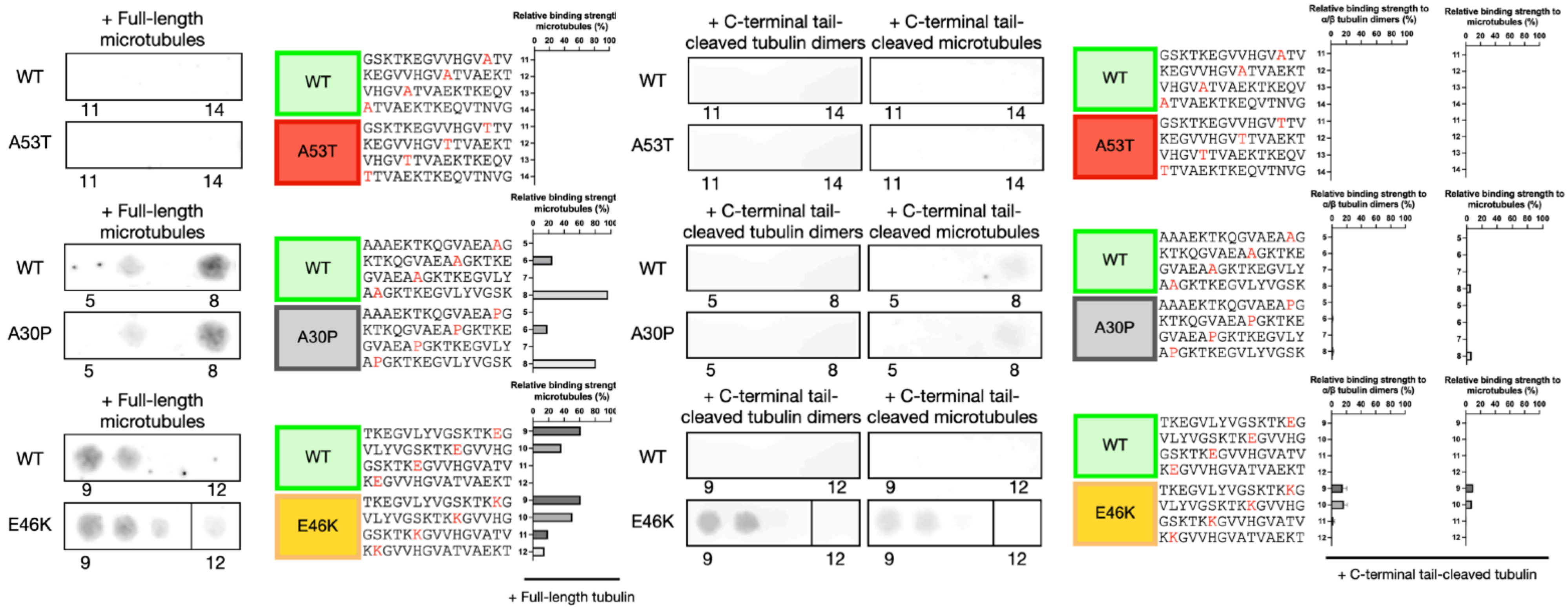


**B** Additional microtubule dynamics parameters and comet density at SYP1⁺ boutons in dopaminergic axons. Panel above: after α-synuclein knockdown and rescue with either ORF-Stuffer control, WT, A53T, A30P, or E46K α-synuclein. Panel below: after infection with shNC and overexpression of either ORF-Stuffer control, WT, A53T, A30P, or E46K α-synuclein.

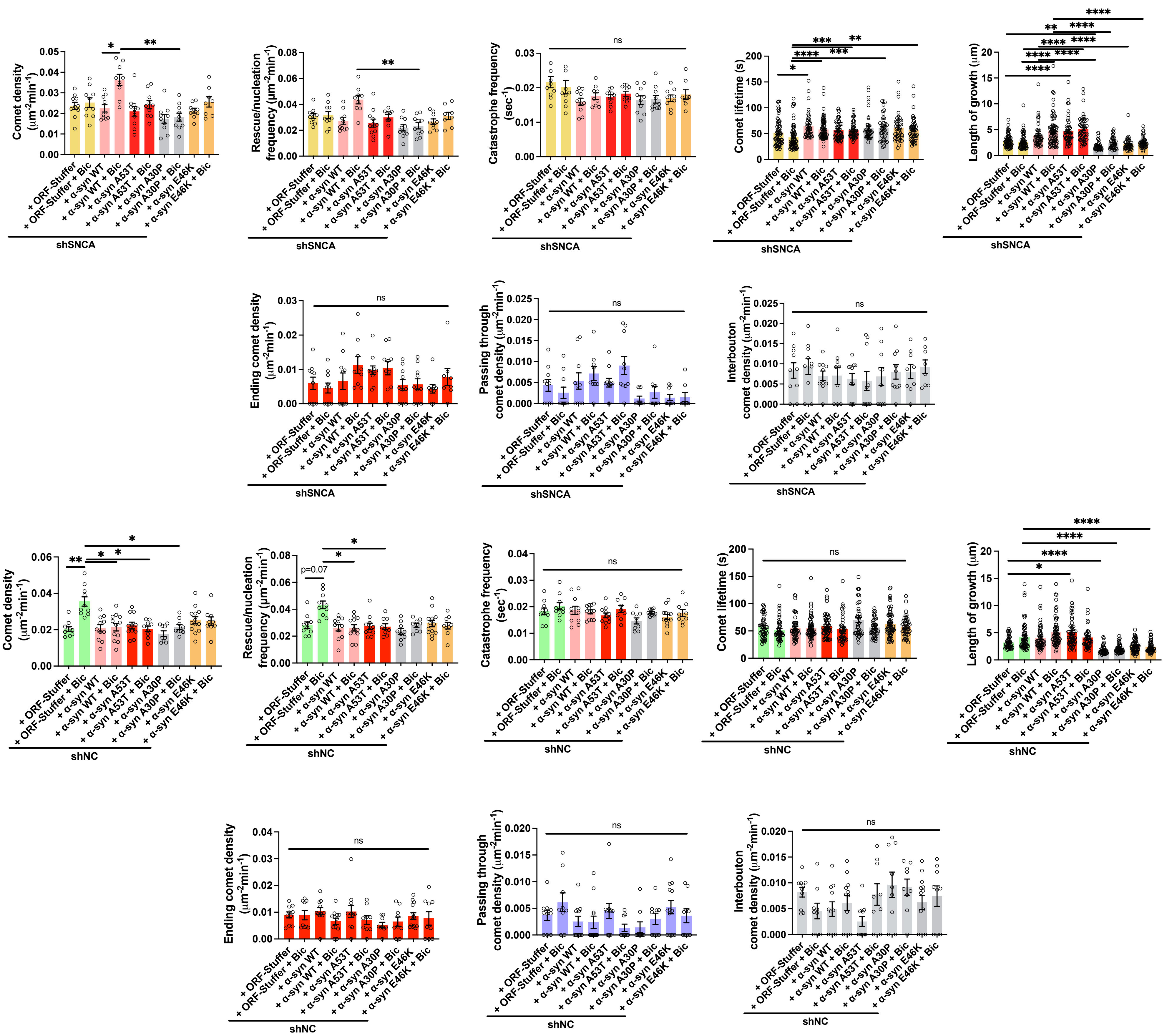

**Figure S1. Microtubule dynamics in dopaminergic neurons under basal conditions and glutamatergic blockade.**

**(A)** Quantification of catastrophe frequency, comet lifetime, and length of growth of EB3 comets in proximal (x<100 µm from cell body) or distal (x>100 µm from cell body) axons and dendrites in DAT-Cre neurons (14-16 DIV). Data are mean ± SEM; ns, p > 0.05; Kruskal-Wallis non-parametric ANOVA. **(B)** Quantification of microtubule dynamics parameters in GCamP3S-DAT-IRES-Cre neurons upon bicuculline stimulation alone or in the presence of NMDA and AMPA receptor blockers to inhibit glutamatergic input. Data are mean ± SEM; *** p<0.001, Kruskal-Wallis non-parametric ANOVA.

**Figure S2. Validation of γ-tubulin depletion, synaptosomal purity, and effects on presynaptic microtubule dynamics in dopaminergic neurons.**

**(A)** Manders colocalization coefficient M1 (γ-tubulin overlapping synapsin1) and M2 (synapsin1 overlapping γ-tubulin) before and after cross-correlation analysis with a 15-pixel X-axis shift applied to each confocal image. Data are mean ± SEM; ****p<0.0001, Mann-Whitney non-parametric test. **(B)** Assessment of crude synaptosomal fraction purity by western blot for GM130 (Golgi marker), histone 2A (nuclear marker), and total protein loading (Ponceau S). **(C)** Quantitative western blot analysis of γ-tubulin levels in dopaminergic neurons (16 DIV) infected for 7 days with non-coding control shRNA (shNC) or shRNA targeting γ-tubulin (shγ-tub). Data are mean ± SEM; ***p<0.001, Mann-Whitney non-parametric test. Live-imaging analysis of bouton density, interbouton distance, and bouton size relative to stable synaptophysin1 (SYP1)$^{+}$ puncta in DAT-Cre neurons (14 DIV) infected for 7 days with non-coding shRNA control (shNC) or shRNA targeting γ-tubulin (shγ-Tub). Data are mean ± SEM; ns, p > 0.05; Mann-Whitney non-parametric test. **(D)** Quantification of comet density, catastrophe frequency, length of growth, growth rate, and comet lifetime in axons of DAT-Cre neurons expressing flexed EB3-mGreenLantern and SYP1-mRuby after 7 days of lentiviral delivery of shRNA targeting γ-tubulin (shγ-Tub), non-coding control (shNC) at baseline or after bicuculline incubation. Data are mean ± SEM; *** p<0.001, Kruskal-Wallis non-parametric ANOVA. Quantification of ending, passing through, and interbouton comet density relative to stable SYP1$^{+}$ puncta in DAT-Cre neurons under the conditions shown in Fig. 2F. Data are mean ± SEM; ns, p > 0.05; Kruskal-Wallis non-parametric ANOVA.

**Figure S3. γ-Tubulin–dependent presynaptic microtubule nucleation supports activity-evoked synaptic vesicle trafficking and dopamine release kinetics.**

**(A)** Quantification of total moving SYP1$^{+}$ puncta, starting SYP1$^{+}$, and ending SYP1$^{+}$ events relative to stable SYP1$^{+}$ puncta, and total anterograde, retrograde, and bidirectional SYP1$^{+}$ tracks in axons of DAT-Cre neurons expressing flexed SYP1-EGFP after 7 days of lentiviral delivery of shRNA targeting γ-tubulin (shγ-Tub), non-coding control (shNC) or acute treatment with the γ-tubulinselective inhibitor GatastatinG2 (Gat) at baseline or after bicuculline incubation (shNC + Bic, shγ-Tub + Bic, and shNC + Gat + Bic) or combined (shγ-Tub + Gat). Data are mean ± SEM; **** p<0.0001, Kruskal-Wallis non-parametric ANOVA. **(B)** Effect of GastastatinG2 on half-decay time ($t_{1/2}$) of evoked DA release in WT and α-synuclein KO slices at a 2- 3-hour incubation window. Each data point represents an individual slice. Pairwise comparisons are shown for control and GastastatinG2-treated slices (N=9 WT and N=10 α-synuclein KO pairs of slices, respectively; ns, p > 0.05, paired t-test). **(C)** Representative FSCV traces of dopamine release evoked by a train of 5 stimuli at 40 Hz (left) and corresponding quantification of pooled dopamine release from WT and α-synuclein KO mice (right). WT N=6; α-synuclein KO N=4; n= at least 90 recording spots. Data are mean ± SEM; ** p<0.01, Mann-Whitney nonparametric test.

**Figure S4. Effects of α-synuclein depletion on γ-tubulin localization, presynaptic structure, vesicle trafficking, and tubulin-binding specificity of α-synuclein.**

**(A)** Quantitative western blot analysis of α-synuclein levels in dopaminergic neurons (16 DIV) infected for 7 days with non-coding control shRNA (shNC) or shRNA targeting α-synuclein (shSNCA). Data are mean ± SEM; **p<0.01, Mann-Whitney non-parametric test. Live-imaging analysis of bouton density, interbouton distance, and bouton size relative to stable $SYP1^{+}$ puncta in DAT-Cre neurons (14 DIV) infected for 7 days with non-coding control shRNA (shNC) or shRNA targeting α-synuclein (shSNCA). Mean ± SEM; ns, p > 0.05; Mann-Whitney non-parametric test. **(B)** Maximum-intensity projection of spinning disk confocal images from wild-type ventral midbrain neurons (16 DIV) fixed and stained for γ-tubulin, TH, and synapsin 1/2 after infection with shNC or shSNCA. White arrows indicate γ-tubulin puncta co-localizing with synapsin 1/2 within a TH mask. Manders coefficients of $synapsin1/2^{+}$ puncta co-localizing with γ-tubulin (M1, fraction of γ-tubulin overlapping synapsin1/2 on the TH mask; M2, fraction of synapsin1/2 + puncta overlapping γ-tubulin on the TH mask) and quantitative γ-tubulin immunofluorescence intensity normalized to TH in dopaminergic neurons treated as in (A). Data are mean ± SEM; ns, p > 0.05; Mann-Whitney non-parametric test. **(C)** Quantification of catastrophe frequency in axons of DAT-Cre neurons expressing flxed EB3-mGreenLantern and SYP1-mRuby after 7 days of lentiviral delivery of shRNA targeting α-synuclein (shSNCA), non-coding control (shNC) at baseline or after bicuculline incubation. Data are mean ± SEM; *** p<0.001, Kruskal-Wallis non-parametric ANOVA. Quantification of interbouton comet density relative to stable $SYP1^{+}$ puncta in DAT-Cre neurons treated as in Fig. 4C. Data are mean ± SEM; ns, p > 0.05; Kruskal-Wallis non-parametric ANOVA. **(D)** Quantification of total moving SYP1, combined starting & ending SYP1 events, starting SYP1, and ending SYP1 relative to stable SYP1+ puncta, and of anterograde, retrograde, and bidirectional SYP1 tracks in axons of DAT-Cre neurons expressing SYP1-mRuby after 7 days of lentiviral delivery of shRNA targeting α-synuclein (shSNCA) or non-coding control (shNC), at baseline or after bicuculline incubation (shNC + Bic, shSNCA + Bic). Data are mean ± SEM; ** p<0.01, Kruskal-Wallis non-parametric ANOVA. **(E)** Manders coefficients for $synapsin1/2^{+}$ puncta overlapping γ-tubulin in dopaminergic neurons treated as in Fig. 4G (M1, fraction of γ-tubulin overlapping synapsin1/2 on the TH mask;

M2, fraction of synapsin1/2$^{+}$ puncta overlapping γ-tubulin on the TH mask), and for synapsin1/2$^{+}$ puncta co-localizing with endogenous α-synuclein on the TH mask.

**Figure S5. α-Synuclein C-terminus controls activity-evoked presynaptic microtubule nucleation at *en passant* boutons in dopaminergic neurons**

**(A)** Representative western blot of purified porcine α/β-tubulin incubated with increasing Subtilisin A: tubulin ratios showing loss of tyrosinated α-tubulin signal at 1:60, indicative of complete cleavage of α-tubulin C-terminal tail. Below, representative pep-spot results from membranes bearing WT α-synuclein peptides incubated with 2.5µg/ml C-terminal tail-cleaved α/β tubulin dimers, C-terminal tail-cleaved polymerized microtubules, or polymerized microtubules assembled from full-length tubulin. Right, sequence of the 33 WT α-synuclein peptides and relative binding strength (%) to C-terminal tail-cleaved α/β tubulin dimers, C-terminal tail-cleaved polymerized microtubules, or polymerized microtubules assembled from full-length tubulin. Data are mean ± SEM; n=2 membranes and two independent preparations for C-terminal tail-cleaved α/β tubulin dimers. n=1 for C-terminal tail-cleaved polymerized microtubules and for polymerized microtubules assembled from full-length tubulin. **(B)** Quantification of rescue/nucleation frequency, catastrophe frequency, length of growth, growth rate, and comet lifetime in axons of DAT-Cre neurons expressing flexed EB3-mGreenLantern and SYP1-mRuby after 7 days of lentiviral delivery of shRNA targeting α-synuclein (shSNCA), at baseline or after bicuculline incubation. α-Synuclein knockdown was rescued with either control (ORF-Stuffer) (yellow), WT α-synuclein (pink) or a C-terminally truncated α-synuclein (1-125) (brown). Data are mean ± SEM; **** $p<0.0001$, Kruskal-Wallis non-parametric ANOVA. Quantification of ending, passing through, and interbouton comet density relative to stable SYP1$^{+}$ puncta in DAT-Cre neurons treated as in (B). Data are mean ± SEM; ns, $p > 0.05$; Kruskal-Wallis non-parametric ANOVA.

**Figure S6. Activity-dependent α-synuclein Ser129 phosphorylation drives presynaptic microtubule initiation at *en passant* boutons in dopaminergic neurons without broadly altering microtubule dynamics or tubulin binding to microtubules.**

**(A)** Left, representative pep spot results for the last four peptides (30-33) of human WT, S129A, and S129D α-synuclein incubated with 2.5µg/ml polymerized microtubules assembled from full-length α/β tubulin, developed after incubation with infrared-conjugated antibodies. Sequences of peptides 30-33 with the position of Ser129 (WT), S129A, or S129D highlighted in red and corresponding binding strength (%) to polymerized microtubules (n=1). Right, representative pep spot results for peptides 30-33 of WT, S129A, and S129D α-synuclein incubated with 2.5µg/ml C-terminal tail-cleaved α/β tubulin dimers or C-terminal tail-cleaved polymerized microtubules, with peptide sequences (S129, S129A, S129D highlighted in red) and relative binding strength (%) to these species. Data are mean ± SEM; n=2 membranes and two independent preparations for C-terminal tail-cleaved α/β tubulin dimers; n=1 for C-terminal tail-cleaved polymerized microtubules. **(B)** Quantitative pS129 α-synuclein immunofluorescence intensity expressed as integrated density (A.U.) calculated on ectopically expressed flex SYP1-mRuby mask in DAT-Cre dopaminergic neurons after 1h incubation with bicuculline or after 2h incubation with PLK2 inhibitor BI 2536 + bicuculline after imaging of experiment panel (C). Data are mean ± SEM; ** $p<0.01$, Kruskal-Wallis non-parametric ANOVA. **(C)** Quantification of comet density, rescue/nucleation frequency, catastrophe frequency, length of growth, growth rate, and comet lifetime in axons of DAT-Cre neurons expressing flexed EB3-mGreenLantern and SYP1-mRuby after 7 days of lentiviral delivery of shRNA targeting α-synuclein (shSNCA), non-coding control (shNC), at baseline or after bicuculline incubation. The shNC condition expressed the control vector (TagBFP2-T2A-ORF-Stuffer) (green) while α-synuclein knockdown was rescued with phospho-mimetic S129D (magenta), or phospho-null S129A α-synuclein (purple). The control condition was also incubated with the PLK2 inhibitor BI 2536 (orange). Data are mean ± SEM; **** $p<0.0001$, Kruskal-Wallis non-parametric ANOVA. Quantification of ending, passing through, and interbouton comet density relative to stable SYP1$^{+}$ puncta in DAT-Cre neurons under the conditions shown in Fig. 6D. Data are mean ± SEM; ns, $p > 0.05$; Kruskal-Wallis non-parametric ANOVA. **(D)** Maximum-intensity projection of spinning disk confocal images from wild-type ventral midbrain neurons (16 DIV) fixed and stained for TH, synapsin 1 and either PLA α-synuclein/γ-tubulin or PLA pS129 α-synuclein/γ-tubulin. Negative control was performed by removing the α-synuclein primary antibody. Below: quantification of synaptic PLA puncta density expressed as puncta/μm2 for PLA α-synuclein/γ-tubulin or PLA pS129 α-synuclein/γ-tubulin with or without a 3 h incubation with bicuculline to stimulate neuronal activity. PLA α-synuclein/γ-tubulin was also compared to the negative control PLA -/γ-tubulin without α-synuclein antibody. Data are mean ± SEM; ****, $p < 0.0001$; Mann-Whitney non-parametric test.

**Figure S7. PD mutants A53T, A30P, and E46K α-synuclein differentially disrupt presynaptic microtubule initiation and growth at *en passant* boutons in dopaminergic neurons.**

**(A)** Left: representative pep spot results for the four peptides mapping each PD mutant (A53T, A30P, and E46K) of α-synuclein incubated with 2.5 μg/ml polymerized microtubules assembled from full-length α/β tubulin, developed after incubation with infrared-conjugated antibodies. Peptide sequences mapping each PD mutant of α-synuclein (A53T, A30P, and E46K), highlighted in red, along with corresponding binding strength (%) to polymerized microtubules assembled from full-length tubulin. Right: representative pep spot results for the four peptides mapping each PD mutant (A53T, A30P, and E46K) of α-synuclein incubated with C-terminal tail-cleaved α/β tubulin dimers or C-terminal tail-cleaved polymerized microtubules. Peptide sequences mapping each PD mutant of α-synuclein (A53T, A30P, and E46K), highlighted in red, along with relative binding strength (%) to C-terminal tail-cleaved α/β tubulin dimers or C-terminal tail-cleaved polymerized microtubules. Data are mean ± SEM; n=2 membranes and two independent preparations for C-terminal tail-cleaved α/β tubulin dimers. n=1 for C-terminal tail-cleaved polymerized microtubules and for polymerized microtubules assembled from full-length tubulin. **(B)** Panel above: quantification of comet density, rescue/nucleation frequency, catastrophe frequency, comet lifetime, and length of growth in axons of DAT-Cre neurons expressing flexed EB3-mGreenLantern and SYP1-mRuby after 7 days of lentiviral delivery of shRNA targeting α-synuclein (shSNCA), at baseline or after bicuculline incubation. α-Synuclein knockdown was rescued with control (ORF-Stuffer) (yellow), WT, A53T (red), A30P (grey), or E46K (orange) α-synuclein. Data are mean ± SEM; **** $p<0.0001$, Kruskal-Wallis non-parametric ANOVA. Quantification of ending, passing through, and interbouton comet density relative to stable $SYP1^{+}$ puncta in DAT-Cre neurons treated as above. Data are mean ± SEM; ns, $p > 0.05$; Kruskal-Wallis non-parametric ANOVA. Panel below: quantification of comet density, rescue/nucleation frequency, catastrophe frequency, comet lifetime, and length of growth in axons of DAT-Cre neurons expressing flexed EB3-mGreenLantern and SYP1-mRuby after 7 days of lentiviral delivery of shRNA control (shNC), at baseline or after bicuculline incubation. Control (ORF-Stuffer) (yellow), WT, A53T (red), A30P (grey), or E46K (orange) α-synuclein were overexpressed alongside endogenous α-synuclein. Data are mean ± SEM; **** $p<0.0001$, Kruskal-Wallis non-parametric ANOVA. Quantification of ending, passing through, and interbouton comet density relative to stable $SYP1^{+}$ puncta in DAT-Cre neurons, treated as above. Data are mean ± SEM; ns, $p > 0.05$; Kruskal-Wallis non-parametric ANOVA.
